# Abundance-occupancy relationships along taxonomic ranks reveal a consistency of niche differentiation in marine bacterioplankton with distinct lifestyles

**DOI:** 10.1101/2021.04.09.439152

**Authors:** Dandan Izabel-Shen, Anna-Lena Höger, Klaus Jürgens

## Abstract

Abundance-occupancy relationships (AORs) are an important determinant of biotic community dynamics and habitat suitability. However, little is known about their role in complex bacterial communities, either within a phylogenetic framework or as a function of niche breadth. Based on data obtained in a field study in the St. Lawrence Estuary, we used 16S rRNA gene sequencing to examine the vertical patterns, strength, and character of AORs for particle-attached and free-living bacterial assemblages. Free-living communities were phylogenetically more diverse than particle-attached communities. The dominant taxa were consistent in terms of their presence/absence but population abundances differed in surface water vs. the cold intermediate layer. Significant, positive AORs characterized all of the surveyed communities across all taxonomic ranks of bacteria, thus demonstrating an ecologically conserved trend for both free-living and particle-attached bacteria. The strength of the AORs was low at the species level but higher at the genus and phylum levels. These results demonstrate that an assessment of the distributions and population densities of finely resolved taxa does not necessarily improve determinations of apparent niche differences in marine bacterioplankton communities at regional scales compared with the information inferred from a broad taxonomic classification.

**Subject Category:** microbial population and community ecology

## Introduction

Aquatic bacterioplankton can either attach to particles or live freely in the water column (Grossart, 2010; Dang and Lovell, 2016). The resulting free-living (FL) and particle-associated (PA) assemblages strongly differ in their community composition and diversity, as shown in several aquatic habitats (e.g., Jackson et al., 2014; Salazar et al., 2015; Milici et al., 2017). For PA assemblages, their community properties depend on the composition of the particles (Simon et al., 2002) and on the particle types (Simon et al., 2002; Rieck et al., 2015; Bižić-Ionescu et al., 2018), which vary strongly from coastal to open waters. Whereas coastal particles mostly originate from riverine inputs, land runoff, and anthropogenic activities (Rieck et al., 2015; Bižić-Ionescu et al., 2018), oceanic particles are more related to biotic processes such as phytoplankton blooms. In addition, empirical studies suggest that FL and PA bacteria differ in their size and lifestyle and respond differently to environmental fluctuations (Dang and Lovell, 2016, and references therein; Adyari et al., 2020). PA bacteria are generally distinguished by their larger genome sizes and larger number of transporters than found in their FL counterparts (Smith et al., 2013). Among the unique genes PA bacteria are those enabling surface colonization and thereby adaption to environmental fluctuations (DeLong et a., 2006). Generally, the relative contributions of FL and PA bacteria to whole-community functions differ as well. Thus, while FL bacteria dominate in terms of biomass, PA bacteria, despite being less numerous, readily utilize organic aggregates and are thus responsible for a large amount of microbial activity and production (Simon et al., 2002; Dang and Lovell, 2016). Given the distinct roles of FL and PA assemblages in driving different community features, an in-depth understanding of the diversity and dynamics of these ecological groups and of their environmental relationships is a prerequisite for predicting their response to environmental change.

To determine the niche breadth of a species, both the abundance and the occurrence of that species in different environments must be known. Abundance-occupancy relationships (AORs), originally developed for use in macroecology (Gusto et al., 2000; Freckleton et al., 2006), reflect the correlation between the number of sites occupied by a species and the average local abundance of individuals of that species at the occupied site. Positive AORs have been determined for a wide range of organisms (e.g., Gaston et al., 2000; Soininen and Heino, 2005; Barberán et al., 2012; Liu et al, 2015; Shade et al., 2018; Grady et al., 2019). Previous studies highlighted the importance of feedback between local abundance and occupancy in the identification of taxa with strong temporal signatures, thus demonstrating the ability of AORs to reveal large-scale microbial community-environment relationships and taxon persistence (Barberán et al., 2012; Choudoir et al., 2017; Grady et al., 2019). In terms of metacommunity dynamics, changes in local abundance and regional distribution may be a function of a species’ niche breadth (Barberán et al., 2012): the broader the niche, the wider the distribution and the greater the local abundance (Lengyel et al., 2020). For microorganisms, even close relatives may be ecologically and physiologically distinct and thereby occupy distinct microbial habitats (Hunt et al., 2008; Koppel and Wu, 2013; Larkin and Martiny, 2017). Analyzing AORs across taxonomic ranks (from the species to the phylum level) may therefore provide insights into the ecological coherence that underlies habitat-taxon relationships.

The relevance of bacterial taxa as biologically meaningful units is widely discussed in microbial ecology, accompanied by investigations of the ecological consistency of higher taxonomic levels (Koeppel et al., 2007; Philippot et al. 2010; Koeppel and Wu, 2012; Fernández et al., 2018). Bacteria assessed at taxonomic ranks above the species level (i.e., genus to phylum) exhibit some ecological coherence, e.g., in their habitat preferences (Philippot et al. 2010) but the phylogenetic depth of that coherence varies among bacterial lineages (Koeppel and Wu, 2012). Moreover, the patterns identified at higher taxonomic ranks may fail to capture the dynamics of the members at lower taxonomic levels (Ruiz-González et al., 2015). Thus, while a broad taxonomic classification may be sufficient to delineate community-environment relationships (Herlemann *et al*., 2016; Lu *et al*., 2016), more finely resolved taxonomic analyses may be needed to predict community dynamics (Needham *et al*., 2017).

St. Lawrence Estuary (SLE), the deepest and largest estuary in the world, is located on the Canadian east coast and connects the St. Lawrence river over a length of 350 km with the Gulf of St. Lawrence, which separates it from the North Atlantic by nearly 1000 km (Vincent and Dodson, 1999). The water column of the SLE is stratified in summer, as the deeper cold water is trapped underneath a newly formed warmer layer at the surface (Saucier, 2003). Microbially mediated biogeochemical processes and taxon-specific metabolic pathways have been well-documented in the SLE (Thibodeau et al., 2010; Bourgoin et al., 2010; Ramachandran and Walsh, 2015) but the catalogue of whole microbial communities that inhabit the estuary have received less attention. A recent study characterized the PA and FL bacterial communities across the Lower SLE (Cui et al., 2020) but little is known about the alpha-diversity or the relationships between the bacterial community and the environment, and therefore the mechanisms that underlie community assembly.

In this study, the 16S rRNA gene diversity and vertical assembly of FL (0.22–3 μm) and PA (3–200 μm) bacterial assemblages inhabiting the SLE were investigated together with their AORs and the coherence of the observed distributions, both at broad- and fine-scale taxonomic classifications. The following three questions were addressed: 1) Are there vertical patterns (surface and cold intermediate layer) of FL and PA bacterioplankton assembly? If so, what is the relationship between community structure and environmental variables? 2) Among PA and FL communities, can habitat generalists (broad occupancy across sampling sites) and specialists (restricted occupancy to few sites) be distinguished? 3) To what extent can AORs serve as a proxy of the ecological coherence of niche differentiation along taxonomic resolutions?

## Materials and methods

### Sampling sites and water collection

Water samples were collected during an expedition (MSM 046) of the research vessel *Maria S. Merian* to the Gulf of St. Lawrence and the lower St. Lawrence River from August 25 to September 4, 2015. The sampling campaign along 2778 km of the SLE was conducted during a cruise that extended from Halifax to St. John, covering the Lower St. Lawrence Estuary (LSLE) and the Gulf of St. Lawrence (GSL). One of the most distinct features of both regions is a cold intermediate layer (CIL) that develops in summer and is renewed by water from different origins (Smith et al., 2006a; Smith et al., 2006b). During the cruise, surface (3 m) and CIL (50–65 m, according to the lowest temperature in the epipelagic zone) samples were collected from nine stations, including determinations of the water chemistry as a measure of environmental heterogeneity (Figure 1A; Supplementary Table S1).

**Figure 1.**
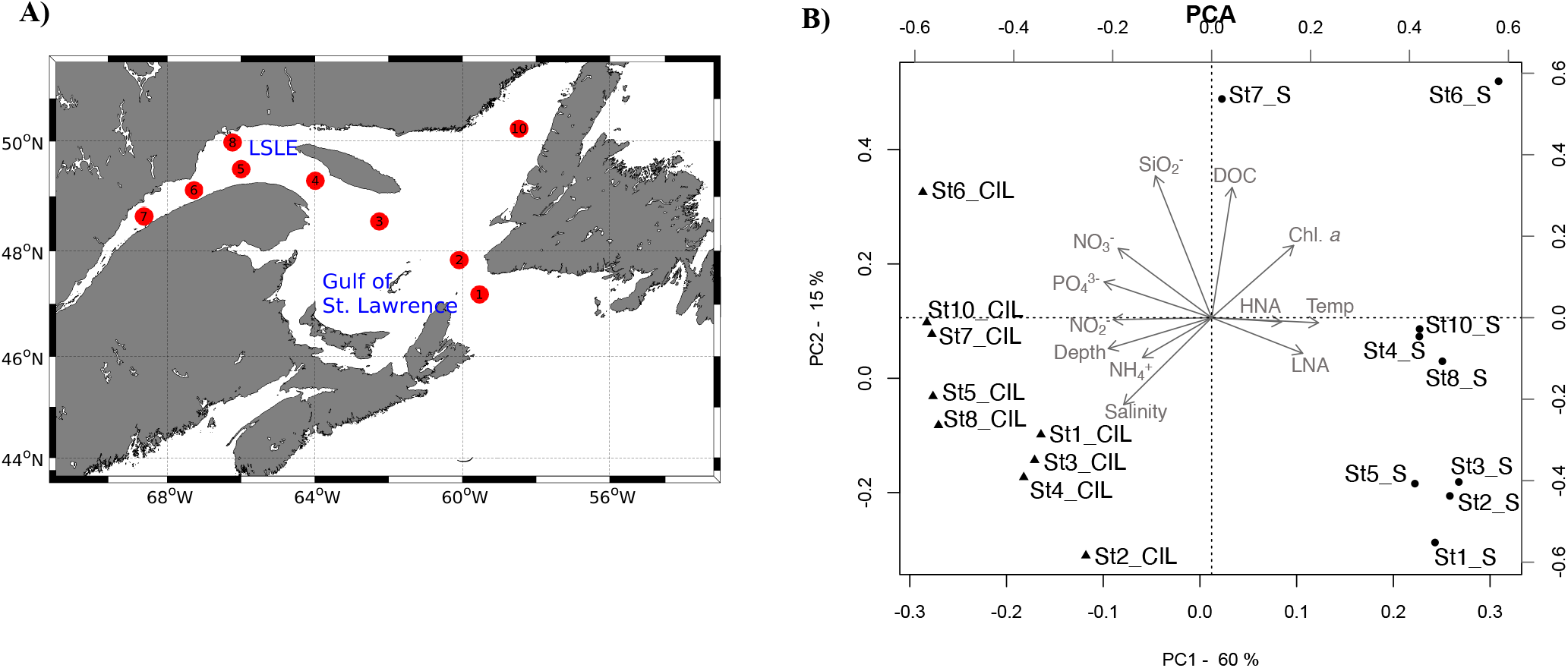
(**A**) Map of the study area, showing the cruise track of the *Maria S. Merian* along the Gulf of St. Lawrence (GSL, stations 1–4) and the lower St. Lawrence Estuary (LSLE, stations 5–8, 10). Sampling stations are labeled consecutively as numbered circles (red), in the order of sampling. (**B**) The results of the principal component analysis (PCA) showing the grouping of the sampling sites in terms of environmental variables. In the plot, sampling sites at the surface (S) are denoted by triangles and those at the cold intermediate layer (CIL) by circles. Gray arrows indicate environmental variables; arrows pointing in the same direction and with a similar length indicate a high-positive correlation with the sampling sites. The station ID and depth of each sampling site are indicated. The two first axes explained 78% of the variance. Abbreviation: DOC: dissolved organic carbon, Temp: temperature, NH4-N: ammonium, NO2-N: nitrite, NO3-N: nitrate, PO4-P: phosphate, SiO2: silica, HNA: bacterial cells with a high nucleic acid content, LNA: bacterial cells with a low nucleic acid content.

### Sample collection and cell partitioning

For each station, water from the two depths was collected using a CTD rosette sampler and filtered through a 200-μm mesh into 20-L carboys to remove large zooplankton. Conductivity, temperature, and chlorophyll-a (Chl-*a*) fluorescence *in situ* at the corresponding depths were recorded using the CTD rosette during acquisition of the water samples. Fifteen mL of the filtered water of each sample was filtered through GF/F filters (Whatman, Daseel, Germany) and then stored at –20°C until used in the nutrient analysis; another 4 mL was preserved with formaldehyde at a final concentration of 2% for the enumeration of bacterial cells by flow cytometry. These measurements were described in detail in a previous study (Shen et al., 2018a). To separate the PA and FL bacterial communities, cells in 1 L of water were partitioned into two size fractions by sequential filtration of the pre-filtered water through 47-mm diameter Durapore filters with mesh sizes of 3.0 µm and 0.22 µm (Millipore, Darmstadt, Germany). The filtration was performed in triplicate for each sample in the same manner. All filters were collected in separate tubes, immediately flash frozen in liquid nitrogen, and stored at –80°C until used for nucleic acid extractions.

### DNA extraction and Illumina amplicon sequencing

DNA was extracted from two of the three replicate filters using the Allprep DNA/RNA mini kit (Qiagen, Hilden, Germany) according to the manufacturer’s protocol and then quantified using a NanoDrop 2000 (Thermo Scientific, Darmstadt, Germany). DNA extraction duplicates were pooled to provide more homogeneous samples, resulting in a total of 36 DNA extracts from distinct water samples. The extracts were PCR-amplified using the bacterial primer pair 341F and 805R (Herlemann et al., 2011). The PCR products were visualized on 1.5% agarose gels before being sent to LGC Genomics GmbH (Berlin, Germany) for high-throughput Illumina sequencing (MiSeq paired-end run with 2 ×300 bp). Each sample was used to obtain duplicate libraries and then sequenced for technical replicates (n = 2). The V3-4 region of the bacterial 16S rRNA gene was sequenced using the primers 341F and 805R (Herlemann et al., 2011).

### Sequence processing and the reproducibility of technical sequencing replicates

The open-sourced expandable bioinformatics software package MOTHUR v.1.36.1 was used to analyze the sequence data obtained from the Illumina MiSeq platform. The MiSeq Standard Operation Protocol (Kozich et al., 2013) was followed but customized to fit the dataset, as described in Shen et al., (2018a). Briefly, the quality-trimmed remaining sequences were aligned with sequences in the SILVA bacterial reference database v.123 (Quast et al., 2013). The resulting sequences were then checked for chimeras using the UCHIME algorithm (Edgar et al., 2011). All sequences recognized as non-bacterial (chloroplast-mitochondria-Archaea-Eukaryota) or that could not be classified were excluded from the dataset. Finally, the sequences were clustered into operational taxonomic units (OTUs) based on a 99% sequence similarity and using the average neighbor method. All OTUs were classified according to the Bayesian classifier (Wang et al., 2007). The most abundant sequence in each OTU was chosen as its representative sequence. OTUs that contained only one or two sequences (singletons or doubletons) were removed from the OTU table. In this study, the OTUs were clustered based on a threshold of ≥ 99% similarity of the 16S rRNA gene sequences, such that the finely resolved microbial taxa approximately corresponded to bacterial “species” (Kim *et al*., 2014).

After quality filtering, the 16S rRNA amplicon dataset generated 2, 645,994 (4, 113, 501 sequences before the omission of single- and doubleton OTUs). Thereafter, 28, 824 OTUs were obtained based on a 99% sequence identity. These OTUs corresponded to 773 genera, 354 families, 177 orders, 94 classes, and 29 phyla. An assessment of the reproducibility of the technical replicates (i.e., sequencing replicates, n=72 in total) for each of the 36 samples showed the similarity among the replicates with respect to their beta-diversity (Supplementary Figure S3). Given the low technical variability, non-normalized technical replicates were pooled into one aggregate set of sequences for each sample to achieve deep coverage of the respective community, by summing up the sequence reads from the technical duplicates. Prior to the downstream analysis, the combined set of sequences was subsampled to 13, 339 sequences (the size of the smallest libraries, see Supplementary Table S1 for details) to standardize the uneven sequencing effort. In addition to our analyses using a 99% sequence similarity for the OTU classifications, we confirmed that the beta-diversity patterns were maintained, indicating the use of exact sequence variants from the DADA2 pipeline (Callahan et al.,2017) for the analyses. The two beta-diversity patterns (Supplementary Figure S1 vs. S2) were found to be statistically indistinguishable based on two separate tests (correlation in a symmetric Procrustes rotation = 0.9711, *p*-value 0.001 on 999 permutations; Mantel correlation = 0.9565, *p*-value 0.001 on 999 permutations). Glassman and Martiny (2018) demonstrated that broadscale ecological patterns are robust in terms of the use of exact sequence variants vs. OTUs. Therefore, strong vertical and size-fractionated patterns can be considered robust regardless of whether 99% OTUs or exact sequence variants are selected. The FASTQ files and associated metadata are publicly available at the European Nucleotide Archive under the accession number PRJEB30352.

### Statistical analyses

#### Environmental relationships among the sites and community diversity

The variability in the environmental variables (inorganic nutrients, Chl *a*, dissolved organic carbon, salinity, temperature, depth, and bacterial abundance) between sampling sites was analyzed by principal component analysis (PCA). Prior to the community analyses, all samples were divided into categorical groups by depth (surface and CIL) and size fraction (FL and PA), resulting in four subcommunities: surface_FL, surface_PA, CIL_FL, and CIL_PA. Within-sample (alpha-) diversity indices, including the observed species richness (S.observed) and evenness (Shannon index/ln S.observed) as well as Faith’s phylogenetic diversity (Faith, 1994), were computed from the subsampled dataset. The observed species richness and evenness were calculated using ‘vegan’ package (Oksanen et al., 2020). For phylogenetic diversity, a phylogenetic tree of all OTUs was constructed using FastTree (Prince et al., 2009), as implemented in QIIME (Caporaso et al., 2010), and used to calculate the sum of the total phylogenetic branch length of species across samples, using the ‘picante’ package. Student’s t-test was used to test the difference in alpha-diversity between the PA and FL communities from the nine stations. Between-sample (beta-) diversity was assessed using the non-multiple dimensional scaling ordination (NMDS) with the Bray-Curtis dissimilarity metric. An analysis of similarity (ANOSIM) was used to test the effects of depth and size fraction on community similarity. A principal coordinates analysis (PCoA) and permutation test were used to analyze the correlation between beta-diversity and environmental factors. Differences in (sub)phylum-level relative abundances between the FL vs. PA communities were determined in a Wilcoxon test.

#### Inferring niche specialization from AORs

The abundance and occupancy patterns of all taxa that made up the subcommunities were analyzed to infer niche specialization patterns in the sampling region. Therefore, two ecological categories with varying degree of niche specialization were defined according to the number of sites at which a taxon was detected (occupancy): taxa present at a minimum of six stations (i.e., an occupancy of >50%) were identified as habitat generalists, and taxa present at a maximum of two stations (i.e., an occupancy of < 25%) as habitat specialists (adopted from Barberán *et al*., 2012).

Once the occupancy of a taxon was determined, the (regional) mean relative abundance of that taxon was calculated by averaging the aggregated relative abundances across the nine stations. Very low mean relative abundances (< 0.002%) were excluded from the analysis, as they were assumed to be highly influenced by sequencing errors (Pandit et al., 2009). This approach was repeatedly used for all taxonomic ranks, i.e., species (99% OTU sequence similarity), genus, family, order, class, and phylum, for the four subcommunities. In addition, Spearman’s rank correlation (*rho*), a non-parametric measure of the degree of association between two variables, was computed to test the correlation between site occupancy and the regional mean relative abundance. The degree of correlation was interpreted as the strength of the AORs and was used in comparisons across taxonomic ranks. *P*-values indicating significant associations were subjected to a Bonferroni correction to account for multiple hypothesis testing.

#### Estimate of changes in niche breadth across taxonomic ranks

The slope of the AORs, representing the rate of change in abundance vs. occupancy, was used to evaluate the degree of niche separation. This analysis was performed for all subcommunities (surface_FL, surface_PA, CIL_FL and CIL_PA) separately. An analysis of the degree of niche separation can offer more information on ecological differences along bacterial taxonomic ranks than provided by phylogeny or taxonomy studies alone (Lu *et al*., 2019). We therefore asked: (1) whether the degree of niche width would be higher at broad rather than at fine taxonomic resolution, as high taxonomic levels comprise subpopulations with divergent niche preferences, and (2) whether the relationship between niche width and taxonomic rank holds true for communities with different lifestyles and origins. Hence, the values obtained from the slope of the AORs at each taxonomic rank, regardless of lifestyle and depth, were used as replicates in an analysis of variance (ANOVA) to test for significant differences (*P* <0.05) in the degree of niche width along taxonomic ranks. To fulfill the ANOVA assumption and determine the normality of the data, a Shapiro-Wilk normality test of the residuals in the linear model was conducted. In the case of significant effects of taxonomic rank on the degree of niche width, Tukey’s *post-hoc* test was used to determine at which level the difference occurred. All statistical analyses and data visualizations were performed in the R environment. The R scripts and computing notes for this study are available on GitHub (https://github.com/IzabelShen/Abundance-occupancy).

## Results

### Water physiochemical characteristics and bacterial abundances

A suite of contextual data was measured for each sampling site, and the relation between environmental variables and the sampling sites was tested (Supplementary Table S1 and Figure 1B). The PCA revealed a clear separation of the surface and CIL in terms of their water physio-chemistry and prokaryotic cell abundance (Figure 1B). The abiotic conditions at the sampling sites were similar at the surface, except at stations 6 and 7, but varied across sites at the CIL. Differences in water temperature, bacterial abundance, and depth had most explanatory values on PC1 (62% variance explained), with contributions from Chl-*a*, nitrite, nitrate, and phosphate. This result suggested their important roles in the differentiation of the two layers. Silica, salinity (and, to a lesser extent, DOC) explained the variation on PC2 (16% variance explained).

### Alpha-diversity

The richness and evenness of the overall communities were greater at the surface than at the CIL (Figure 2A). Faith’s phylogenetic diversity was significantly higher in the FL than in the PA assemblages across stations (Welch’s t-tests, *P*<0.001 for surface; *P*<0.1 for CIL). At the surface, the species richness of PA was marginally higher than that of FL (*P*<0.1), while there was no significant difference in the richness of the two groups at the CIL. Conversely, Pielou’s evenness was higher for FL than for PA assemblages at the CIL (*P*<0.05), suggesting the presence of a small number of highly dominant OTUs in the FL groups of the CIL.

**Figure 2.**
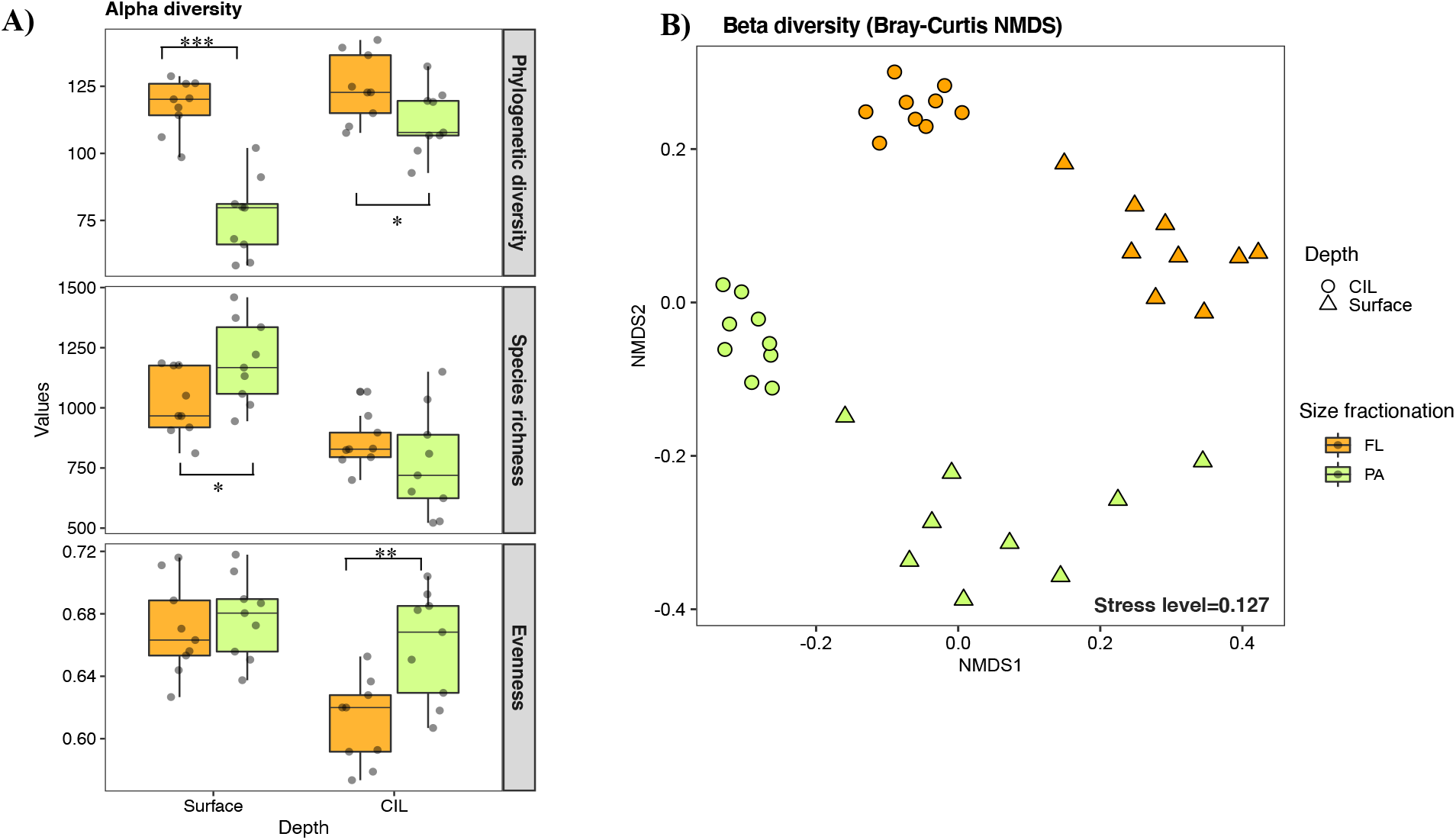
Within-sample (alpha-) (**A**) and between-sample (beta-) (**B**) diversity of the bacterioplankton assemblages inhabiting the St. Lawrence Estuary. The surveyed bacterial communities are defined by depth (surface or cold intermediate layer: CIL) and lifestyle as determined by size-fractionation (free-living: FL indicated by orange or particle-attached: PA indicated by green). For alpha-diversity, the phylogenetic diversity, (realized) species richness, and evenness of PA and FL communities were compared at the surface and the CIL. The significance levels as determined in the corresponding Welch’s t-tests are shown in (**A**): *P*<0.01 ***, *P*<0.05 **, *P*<0.1 *.

### Beta-diversity and divergent community composition

Surface and CIL waters formed diverging environments, as indicated by the PCA (Supplementary Figure S2); hence, taxonomically distinct community compositions were expected for these habitats. Beta-diversity differed between the two water layers in terms of the taxonomic, weighted resemblance (Bray-Curtis dissimilarity metric, Figure 2B). The ANOSIM revealed the significant separation of the four subcommunities from one another by depth (R=0.588, *P*=0.001) and by size fractionation (R=0.671, *P*=0.001; Figure 2B). Independent of their partitioning as FL or PA, bacterial communities inhabiting the CIL were more tightly clustered than those of the surface, suggesting a greater similarity of CIL communities and less similarity between surface-dwelling communities across stations, especially for PA communities. Salinity, silica levels and temperature significantly correlated with FL assemblages at both surface and CIL (PCoA test, *P* <0.01 for all), whereas temperature only correlated with surface-dwelling FL (Table 1, *P*<0.01). The levels of inorganic nutrients (NO_3_^-^ and PO_4_^3-^) correlated significantly with the CIL-dwelling FL assemblages (Table 1) but the correlations between inorganic nutrients and DOC and PA assemblages were weaker. Cells with a low nucleic acid content significantly correlated with surface-dwelling PA assemblages (Table 1).

**Table 1.**
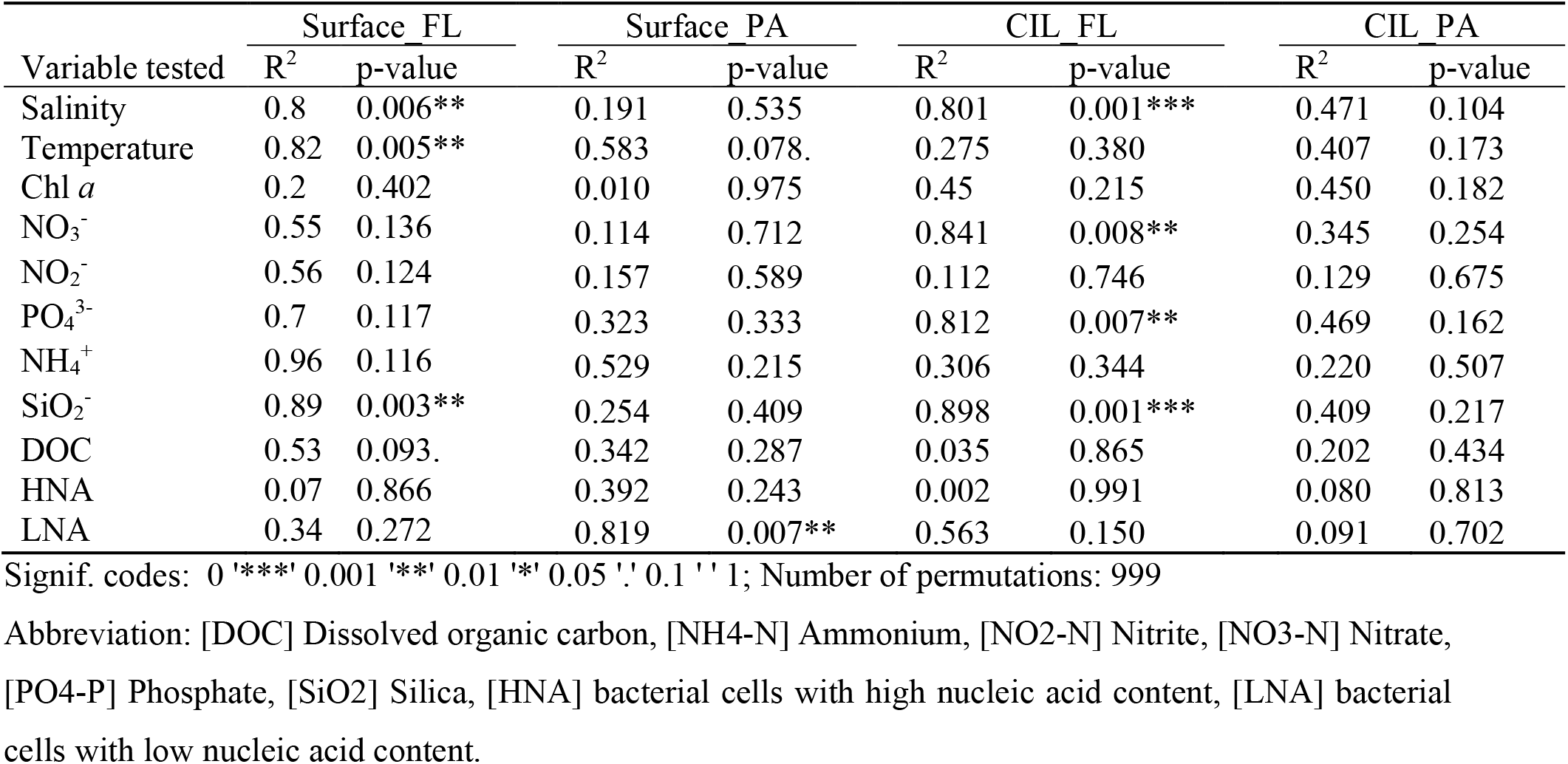
Principal coordinate analysis (PCoA) and permutation test analyzing the correlation between environmental variables and beta diversity. Individual communities surveyed are defined by depth (surface or cold intermediate layer: CIL) and size-fractionation (free living: FL or particle-attached: PA).

An analysis of the taxonomic composition at the phylum/class level revealed that the relative abundances of the dominant taxa were more variable at surface than at CIL stations (Figure 3). Proteobacteria (several subclasses), Bacteroidetes, Actinobacteria, Verrucomicrobia, Cyanobacteria, and Planctomycetes were the dominant bacterial phyla across all stations (Figure 3), with Gammaproteobacteria and Deltaproteobacteria predominating at the CIL and Alphaproteobacteria and Actinobacteria in the surface water. Members affiliated with Lentisphaerae, Chloroflexi, and Deferribacteres were mainly detected at the CIL albeit with relatively low abundances (Figure 3).

**Figure 3.**
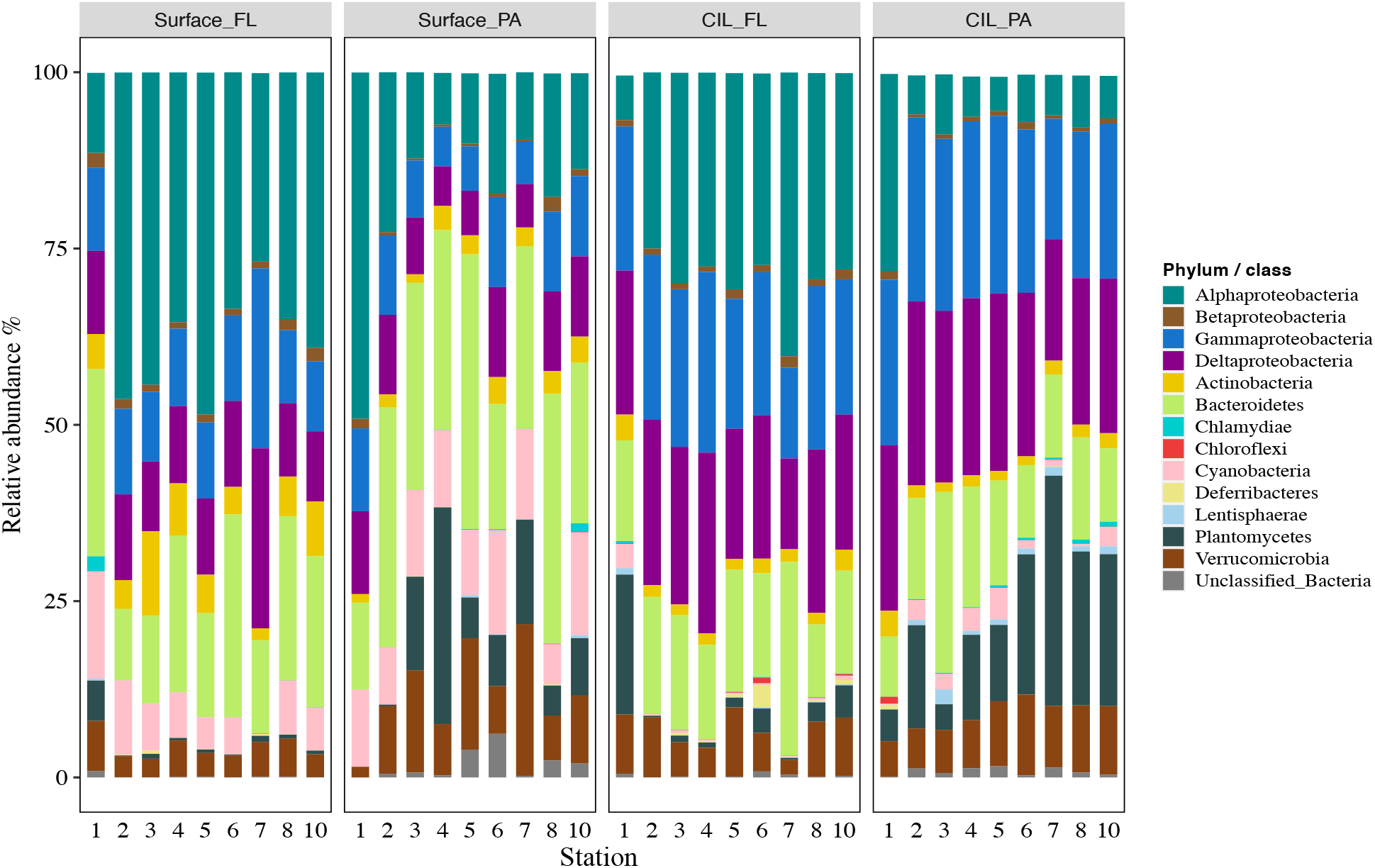
The succession of the dominant bacterial phyla (i.e., collectively >1% relative abundance across samples) of the St. Lawrence Estuary. The mean relative abundances of the (sub-)phyla within the FL and PA communities at two depths, the surface and the cold intermediate layer (CIL) (Surface_FL, Surface_PA, CIL_FL, and CIL_PA), are shown. ‘Unidentified_Bacteria’ represent OTUs that could not be assigned at the phylum level. The phylum Proteobacteria is divided into Alpha-, Beta-, Delta-, Epsilon-, and Gamma-proteobacteria.

Differences in relative abundance among phyla were determined in comparisons between the size-fractionated categories. For example, the relative abundances of members affiliated with Bacteroidetes, Verrucomicrobia, Planctomycetes, and Cyanobacteria were significantly higher in PA than in FL communities (for Bacteroidetes and Verrucomicrobia significant only at the surface) (Wilcoxon test, *P*<0.05; Supplementary Table S2). By contrast, the relative abundances of members affiliated with Actinobacteria, Alphaproteobacteria, Betaproteobacteria, and Deltaproteobacteria were significantly higher in FL communities (Wilcoxon test, *P*<0.05). Despite their overall low contribution, also members affiliated with Deferribacteres, Firmicutes, and Epsilonproteobacteria differed significantly in relative abundance between PA and FL communities (Supplementary Table S2).

### AORs across taxonomic ranks and niche specialization

In addition to community succession, we explored the AORs of the four subcommunities to obtain insights into niche specialization patterns across taxonomic ranks. The results showed a clear trend in the proportions of the two ecological categories along the taxonomic scale (Figure 4:class level; Supplementary Figure S3: other taxonomic levels and Table S3). The number of generalists increased with increasing phylogenetic distance, whereas the opposite was determined for specialists. Additionally, the increase in generalists was larger for PA than for FL communities. This pattern was inverted for specialists, with a more pronounced decrease for FL than for PA communities (Supplementary Table S3). Regardless of the taxonomic rank, the AORs were positive, which suggested that species with low abundances tended to have narrower occupancies and those with high abundances wider occupancies. (Figure 4; Supplementary Table S4). Moreover, the taxonomic composition of specialists vs. generalists differed profoundly for PA and FL bacteria (Supplementary Figure S4). The compositional changes of generalists at the class level generally resembled those at the phylum level across depths (Supplementary Figure S4).

**Figure 4.**
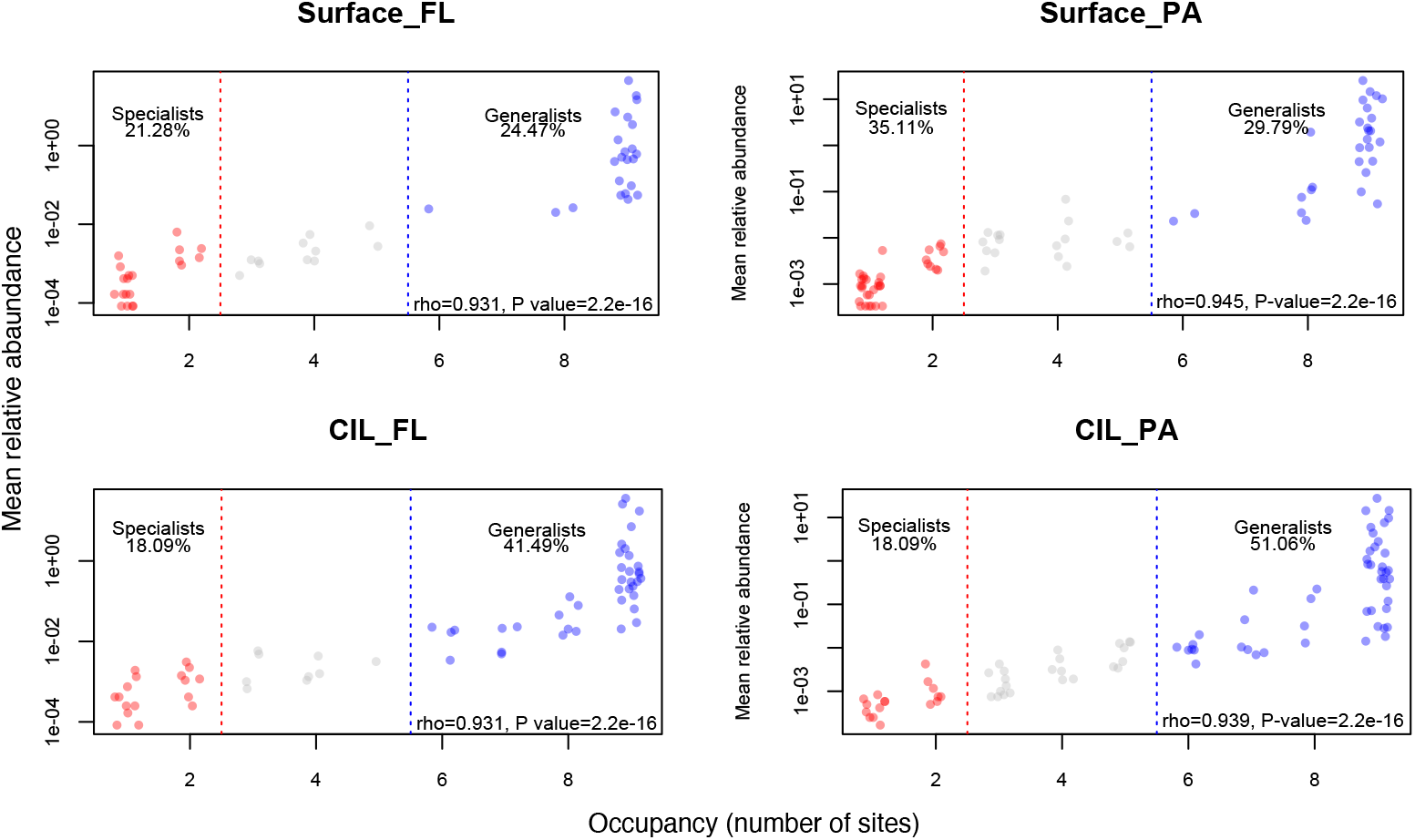
Frequency of occurrence plotted against the mean relative abundance at the class level. The number of occupied sites was used to define the niche breadth of bacteria. Taxa that occupied fewer than two stations were defined as habitat specialists (red), and those occupying more than five stations as habitat generalists. The proportion of the two ecological groups is also shown. The correlation coefficient and *P* value obtained in the Spearman correlation analysis are presented in the lower right corner of each panel.

The strength of the AORs of the PA and FL communities across taxonomic ranks was also considered. We hypothesized that the apparent niche differences of natural bacterial communities would be more clearly defined by the distributions and population densities of finely resolved taxa than by inferences based on a broadly resolved taxonomic classification. Thus, AOR strength was expected to decrease when a broad taxonomic classification was used. However, while the AORs of the surveyed bacterial communities were positive across all taxonomic ranks (Figure 5, Spearman coefficient >0), contrary to our expectation, AOR strength was lowest at the species level and increased at broader taxonomic levels (Spearman coefficient: 0.626–0.719), remaining more or less constant from the genus to the phylum level (Figure 5).

**Figure 5.**
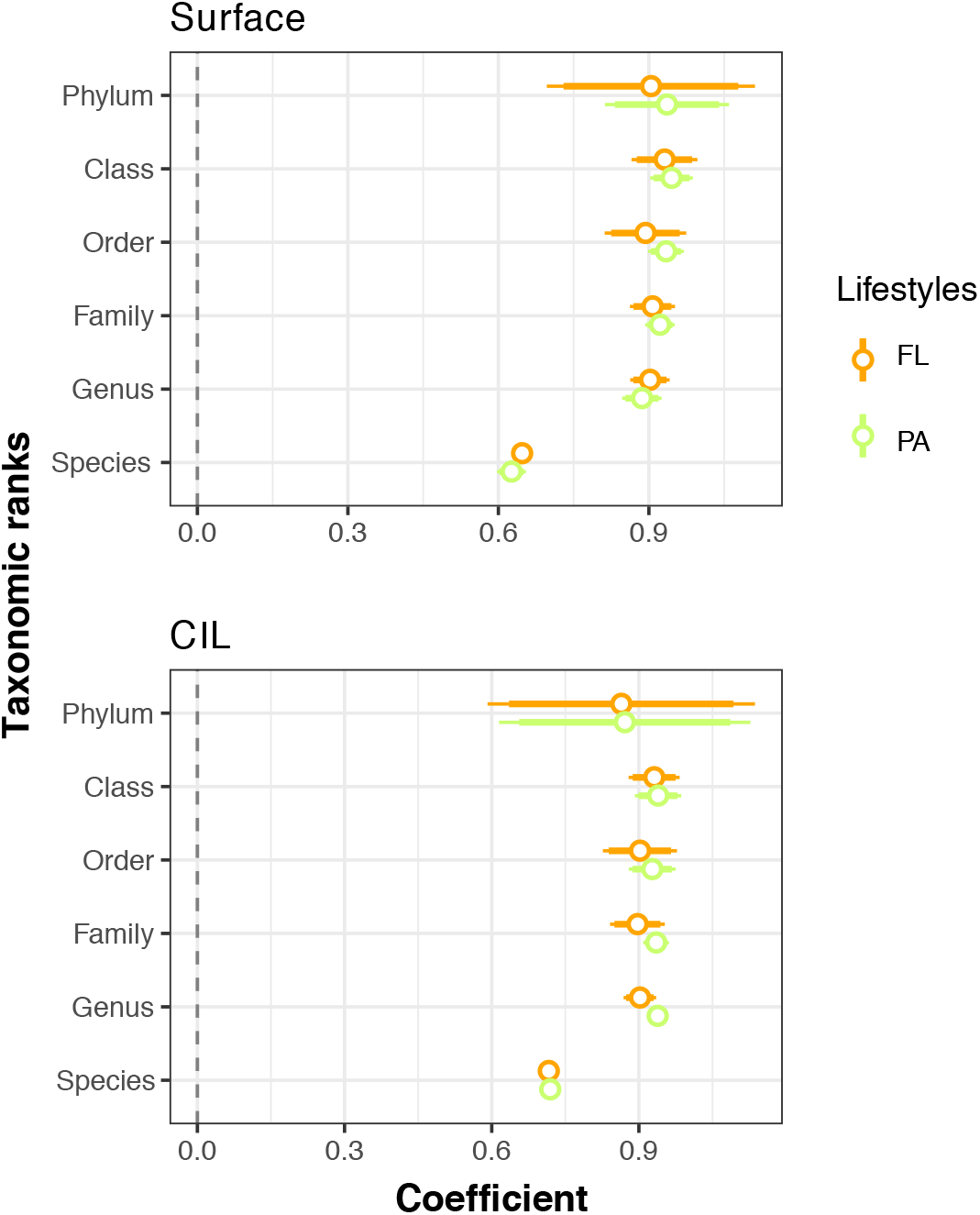
Coefficient plots of the Spearman ranks explaining the mean relative abundance and occupancy relationships (AORs) across taxonomic ranks for free-living (FL, orange) and particle-attached (PA, green) bacterial communities at two depths: the surface and the cold intermediate layer (CIL). The coefficients and their intervals are reported as one and two standard errors of the coefficient. A positive (>0) coefficient indicates a positive correlation of the mean relative abundance of individual taxa with the number of sampled sites at which they occurred (*P*<0.001 in all cases after corrections for multiple testing). The coefficients and *P*-values of the significance for each Spearman’s rank correlation are given in Supplementary Figure S2.

### Degree of niche separation inferred from AORs

Both the slope of the AORs and the taxonomic resolution were analyzed to determine whether the degree of niche separation of the four subcommunities increased with taxonomic rank. The degree of niche separation of the subcommunities increased exponentially with broadening taxonomic resolution (Figure 6, ANOVA: *P* =2.42×10^−6^); thus, increases in the phylogenetic distance resulted in exponential increases in the niche width of the communities. The deviation between communities began at the genus level and became progressively larger until the phylum level, with a significantly large difference at the phylum or class level (Figure 6, inset; Tukey’s *post-hoc, P*<0.001). The increase was quantified by calculating the fold change between every two consecutive taxonomic ranks (Supplementary Table S5B), which again showed that the largest change in niche width occurred between the class and phylum levels. This pattern repeated across four subcommunities with a similar magnitude. Furthermore, the niche index of FL communities deviated from that of the PA communities from the species to the order level (Figure 6).

**Figure 6.**
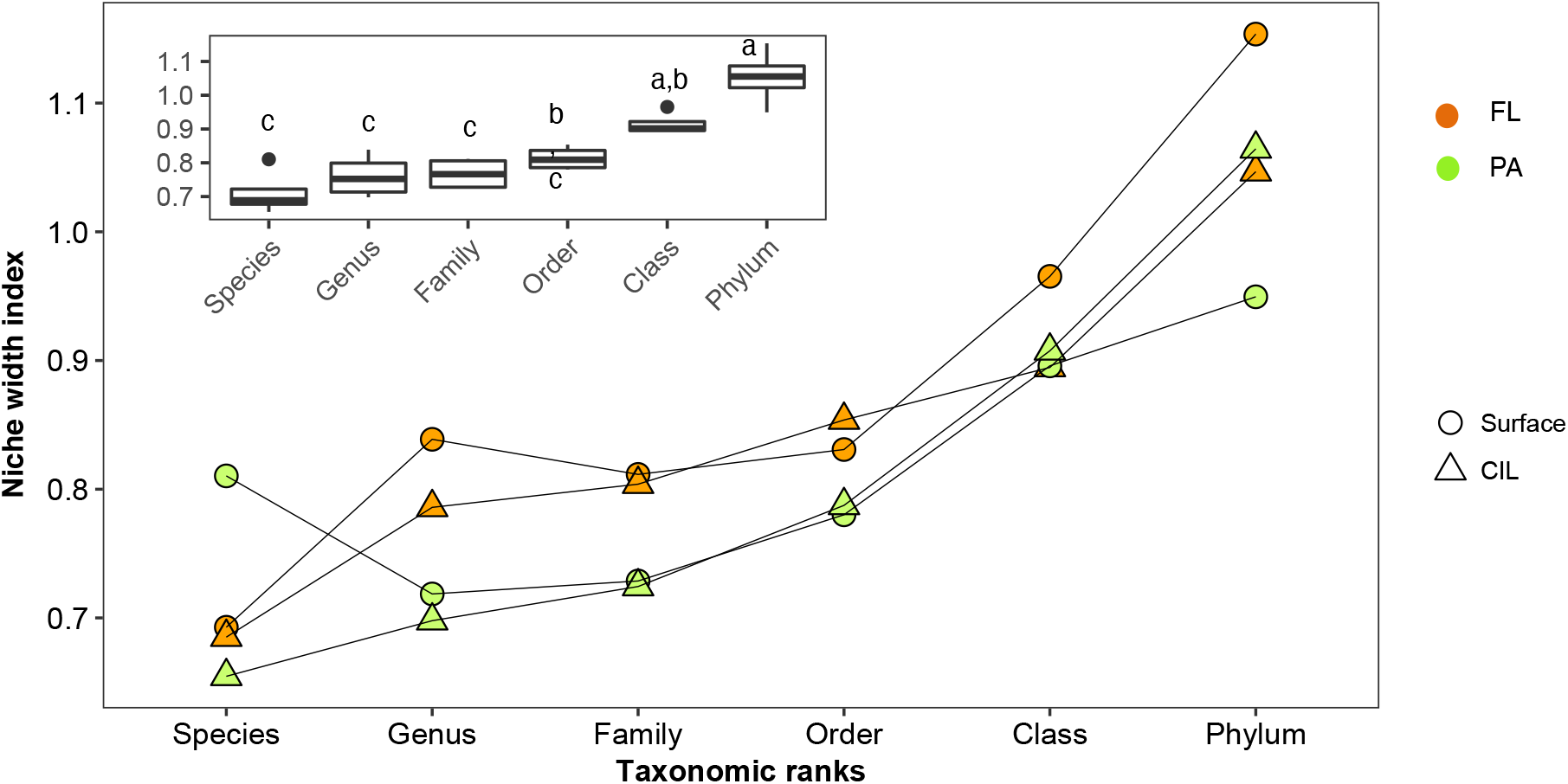
Niche width index (derived from the slope of the logarithm of the abundance values plotted against the number of occupied sites) of the subcommunities along hierarchical taxonomic ranks and the corresponding median rate at each taxonomic rank (inset), without evolutionary information on the bacteria. The different colors indicate the different lifestyles, and the different symbols the different water depths. An analysis of variance (ANOVA) was used to test for significant differences (*P*<0.05) in the niche width index as a function of the taxonomic level. The subcommunities function as replicates for the ANOVA; thus, the values obtained from the subcommunities were used to calculate the variance of these replicates. Tukey’s *post-hoc* test was used to determine the taxonomic level at which the differences occurred. The corresponding homogeneous groups are indicated by a, b and c. The values of the slopes of the AORs of the individual communities are given in Supplementary Table S5A.

## Discussion

### Vertical shifts in community characteristics

A strong stratification of the SLE, with pronounced abiotic differences between the surface and the CIL, was determined during our sampling campaign, consistent with previous reports (Vicent & Dodson, 1999; Elliott & McLusky, 2002). Inorganic nutrient concentrations were higher at the sampling sites of the LSLE than at those located in the GSL, attributable to the nutrient-rich water upwelled and adverted from upstream regions (Therriault & Levasseur, 1985) and to the frequent tidal events in the LSLE (Winkler et al., 2003; Dinauer and Mucci, 2017). Bacterial species richness and evenness were higher at the surface than in the CIL (Figure 2A). The low level of community diversity at the CIL was likely due to the near-freezing water temperature, given that low temperatures suppress the activity and richness of bacterial communities (Kellogg & Deming, 2009).

Community composition was more similar among CIL stations than among surface stations. On the horizontal scale, the conditions in the deeper layer were more similar across stations than that at the surface (Figure 1B). Conversely, there was more heterogeneity in surface water most likely due to phytoplankton bloom and thereby primary production in summer. Dissolved organic carbon released from phytoplankton by direct excretion or through trophic interactions sustains the growth of diverse bacterial assemblages in the surface water (Azam et al., 1983). On a vertical scale, dispersion between the surface and CIL is limited, including vertical mixing not only of the respective environmental matrices but also of microbial cells from each one. It was therefore unlikely that bacteria inhabiting the surface water were able to colonize the CIL.

### Distinct community composition of FL and PA bacteria

The higher phylogenetic diversity of FL than PA communities suggested that the former harbor more phylogenetically diverse groups of bacteria (Figure 2A). However, the species richness and evenness of PA were greater than that of FL at the surface and CIL, respectively (Figure 2A). These results are supported by some (Crespo et al., 2013; Mohit et al., 2014; Rieck et al., 2015) but not all studies, with most of the latter reporting a higher diversity for FL bacteria (Acinas, 1999; Ghiglione et al., 2009; Kellogg & Deming, 2009). The inconsistence across studies can be explained by the environmentally (locality-) dependent origin and material composition of organic particles (Simon et al., 2002; Mohit et al., 2014; Bižić-Ionescu et al., 2018). The strong effects of the measured environmental variables on FL but not on PA assemblages (Table 1) indicated differences in the environmental interactions of species from different lifestyle groups. Thus, PA bacteria may have been influenced by environmental factors that were not assessed in this study, such as the concentrations of particulate organic matter (Mohit, 2014). Estuarine systems are characterized by large inflows of terrestrial materials and allochthonous organic matter (Dinauer and Mucci, 2017), which may have contributed to the high bacterial richness of the PA fraction in the SLE.

In accordance with earlier findings in marine and estuarine systems (Crump et al., 1999; Salazar et al., 2015; Mestre et al., 2017; Cui et al., 2020), the composition of FL bacteria was both taxonomically and phylogenetically distinct from that of PA bacteria (Figure 2B; Supplementary Figure S1). Bacteroidetes, Planctomycetes, and Verrucomicrobia accounted for a larger proportion of the PA fraction than of the FL fraction. These groups have been described previously as PA indicators (Salazar et al., 2015). For example, a strong particle affinity of members of Planctomycetes was reported for biofilm-forming communities on microplastic particles (Wiegand et al., 2020), in the low-saline communities of the Baltic Sea (Rieck et al., 2015), and in deep-sea prokaryotic assemblages (Salazar et al., 2015). In our study, Cyanobacteria were more abundant in the PA than in the FL fractions at all stations, consistent with previous reports of abundant cyanobacterial populations in the LSLE (Cui et al., 2020) but also in the GSL, where the salinity is higher. The presence of cyanobacterial populations on (sinking) particles was also described (Farnelid et al., 2018; Yang et al., 2019). In our study, the high abundance of PA cyanobacteria in the surface layer and in the CIL points to potential bloom conditions in the SLE and thus the potential export of organic materials from the surface to deeper-water habitats via vertical dispersion in the form of particle attachment. Conversely, the high abundances of Alphaproteobacteria and Actinobacteria in the FL fraction in surface water (Figure 3) were consistent with the preferentially pelagic and FL lifestyle of these taxa (Dang and Lovell, 2016).

### AORs and habitat specialization

By applying an established macroecological approach (Gaston et al., 2000) to consider both the occupancy and abundance patterns of bacterial assemblages inhabiting the SLE, we were able to discern two ecological groups: (1) habitat generalists, which tended to be regionally abundant, and (2) habitat specialists, which were regionally rare in all cases, regardless of the taxonomic depth and size fractionation of the bacterioplankton (Supplementary Figure S4). However, data obtained during field and experimental studies demonstrated that, in bacterial communities, habitat specialists were more locally abundant than generalists (Barberán et al., 2012; Logres *et al*., 2013; Shen *et al*., 2018b). The discrepancy can be explained by difference in habitat differentiation, as SLE habitats were less well-differentiated than those surveyed in previous studies of environmental gradients, such that habitat filtering in the SLE was not strong enough to promote the growth of specialists with a high growth rate. These results suggest that definitions of specialists and generalists based on habitat occupancy will depend on the environmental conditions of the sampling area. It has also been suggested that habitat specialists tend to outnumber generalists in ecological communities (Guo *et al*., 2000; Van der Gast *et al*., 2011). However, at least in our study, this tendency skewed towards high taxonomic ranks. Among communities above the order level, generalists made up a larger proportion than specialists (Supplementary Figure S3E, F), with the most pronounced difference occurring at the phylum level. Higher taxonomic ranks are made up of subgroups characterized by increased phylogenetic distances and therefore possibly divergent ecological preferences (Philippot et al., 2010). Therefore, niche breadth will vary as a function of taxonomic resolution, and the niche breadth of individuals should therefore be smaller than that of the species or genus to which they belong. Our results demonstrate that the number and taxonomic resolution of habitat generalists vs. specialists should be taken into account in studies of community dynamics and habitat suitability.

In addition, a significant positive relationship was determined between the mean relative abundance and the mean relative occupancy for the subcommunities surveyed at all taxonomic levels (Supplementary Figure S4). Similar relationships were identified both in lake (Liu et al., 2015) and gut microbial communities (Burns et al., 2016). The AORs were stronger at the class level and weaker at the species level (OTUs at 99% sequence similarity). The high probability of randomness at finer taxonomic scales, such as the species level, may obscure community-environment relationships (Lu et al., 2016).

### Ecological coherence of niche width in FL and PA assemblages

Our data revealed an exponential increase in niche width index (a proxy of ecological niche) with increasing phylogenetic distance, with the sharpest increase occurring between class and phylum (Figure 6; Supplementary Table S5 B). This indicates that bacterial taxa possess an ecological coherence that weakens at higher taxonomic ranks. Philippot et al. (2010) came to a similar conclusion in their investigation of the habitat/phylotype association of a set of microbial strains whose genomes had been sequenced. In contrast to our study, the authors used an approach focused on habitat similarity rather than the divergences of habitats and niche width. Ultimately, they found a negative correlation between habitat similarity and taxonomic ranks and thus reached the same conclusion as resulting from our study, i.e., that the ecological consistency of this relation diminishes above the class level. At higher taxonomic ranks (i.e., class or even phylum level), the niche indices of the subcommunities varied as a function of water depth rather than lifestyle, observed at finer taxonomic scales. These findings suggest that niche differentiation is a phylogenetically conserved tendency of both PA and FL bacteria, as noted elsewhere (Mestre *et al*., 2017), and that the effect of water depth on the entire communities within each lifestyle is stronger when broad taxonomic scales are applied.

### Caveats and recommendations for future work

Our study had several limitations. First, a 99% sequence similarity cutoff was used to aggregate OTUs at the species level, although a cutoff of 97% is more common. This might have led to an overestimation of species, given the inherent sequencing errors and biases. Hence, the OTU-based classification should be interpreted as describing closely related bacterial lineages rather than species. Second, FL and PA bacteria were partitioned based on two cell size categories (0.2–3 µm and >3 µm). Since there is no consensus on how to most effectively separate these groups, caution is needed in comparing the community patterns identified in our study vs. in previous studies, in which different size-fractionation methods were used to partition FL and PA bacteria. Nevertheless, the methods chosen for this study are commonly used in microbial ecology and they provided insights into community-environment relationships in the SLE and the vertical assembly of bacterial communities that differ in their lifestyles.

## Conclusions

Our study showed that the FL fraction consisted of phylogenetically more diverse bacterial populations than the PA fraction, but vertical patterns were detected within each fraction. The significant and positive AORs identified across all taxonomic ranks of bacteria demonstrated an ecologically conserved trend for both lifestyles, even at the small regional scale of the study. AOR strength was low at the species level but increased steadily with progressively higher taxonomic rank, irrespective of bacterial lifestyle. Thus, in studies of bacterial assemblages at regional scales, analyses of high taxonomic ranks may already provide a reasonable understanding of contemporary patterns of their abundance and occurrence. Finally, our analysis of the degree of niche width inferred from the AOR slopes along taxonomic ranks revealed the consistent niche differentiation of bacterioplankton with different lifestyles, with greater differentiations at higher (class and phylum) taxonomic levels. Collectively, our results corroborate previous work (e.g., Shade et al., 2018) in that they extend the principles developed for plants and animals to an understanding of the dynamics and distribution of complex natural microbial communities. Thus, not only is a FL or PA lifestyle a phylogenetically conserved trait for pelagic marine bacterioplankton (Salazar et al., 2015), but the niche differences and distributions of these two groups at regional scales are maintained over broad taxonomic resolution. Future work should ascertain whether these patterns also describe other microbial communities classified by multiple size-fractionations and whether the relationship is scale-dependent.

## Acknowledgments

We thank the captain and crew of MSM cruise 046 for facilitating water collection, Christian Burmeister for inorganic nutrient measurements, Jenny Jeschek for measuring dissolved organic carbon, and Ronny Baaske for the flow cytometric determinations. We also are grateful to Jen-ping Peng for generating a sampling map of the St. Lawrence estuary for this study. The work was supported by a grant (JU 367/15-1) from the German Research Foundation (DFG) to KJ.

## Author contributions

DIS and KJ conceived the study. DIS conducted field and laboratory work, data analysis, and wrote a first version of the manuscript. AL processed samples and performed data analysis. KJ financed the study. All authors discussed the results and commented on the manuscript.

