## Supplemental_Materials for "Abundance-occupancy relationships along taxonomic ranks reveal a consistency of niche differentiation in marine bacterioplankton with distinct lifestyles"

**Supplementary Table S1** Summary of the geolocation, nutrient content, bacterial cell abundances and number of sequence reads of each station. ‘Sample_ID’ represents the sampling station corresponding to the map (e.g., st1), size fractionation (FL, free-living or PA, particle-associated) and depth (S, surface or CIL, cold intermediate layer). Abbreviation: Temp, temperature; DD, decimal degrees; No. seqs, number of sequences; Tech_rep1 and Tech_rep 2: technical sequencing replicate 1 and replicate 2, respectively; NA: no data available.

| Sample_ID | Latitude DD | Longitude DD | Depth m | Tech_rep 1 No. seqs | Tech_rep 2 No. seqs | Aggregated No. Seqs | Salinity | Temp °C | Chl A µM / ml | $NO_3^-$ µM / ml | $NO_2^-$ µM / ml | $PO_4^{3-}$ µM / ml | $NH_4^+$ µM / ml | $SiO_2^-$ µM / ml | DOC µM / ml | bacteria cells / ml |
|---|---|---|---|---|---|---|---|---|---|---|---|---|---|---|---|---|
| St1_FL_CIL | 47.19 | -59.54 | 65 | 32, 137 | 27, 428 | 59, 565 | 32.4 | 1 | 0.056 | 4.713 | 0.161 | 0.705 | < 0.5 | 3.490 | 8.7 | 3.54E+05 |
| St1_FL_S | 47.19 | -59.54 | 3 | 77, 432 | 51, 744 | 129, 176 | 32.9 | 18 | NA | NA | NA | NA | < 0.5 | NA | NA | 4.01E+06 |
| St1_PA_CIL | 47.19 | -59.54 | 65 | 23,476 | 25, 875 | 49, 351 | 32.4 | 1 | 0.056 | 4.713 | 0.161 | 0.705 | < 0.5 | 3.490 | 8.7 | 3.54E+05 |
| St1_PA_S | 47.19 | -59.54 | 3 | 16, 954 | 53, 021 | 69, 975 | 32.9 | 18 | NA | NA | NA | NA | < 0.5 | NA | NA | 4.01E+06 |
| St2_FL_CIL | 47.83 | -60.08 | 65 | 59, 113 | 62, 581 | 121, 694 | 32 | 0.44 | 0.147 | 2.796 | 0.121 | 0.716 | 1.247 | 0.691 | 4.9 | 2.79E+05 |
| St2_FL_S | 47.83 | -60.08 | 2 | 11, 5204 | 59, 537 | 174, 741 | 30.2 | 18.7 | 0.495 | 0.385 | < 0.05 | < 0.1 | < 0.5 | 1.685 | 6.7 | 3.90E+06 |
| St2_PA_CIL | 47.83 | -60.08 | 65 | 30, 902 | 15, 028 | 45, 930 | 32 | 0.44 | 0.147 | 2.796 | 0.121 | 0.716 | 1.247 | 0.691 | 4.9 | 2.79E+05 |
| St2_PA_S | 47.83 | -60.08 | 2 | 42, 313 | 30, 994 | 73, 307 | 30.2 | 18.7 | 0.495 | 0.385 | < 0.05 | < 0,1 | < 0.5 | 1.685 | 6.7 | 3.90E+06 |
| St3_FL_CIL | 48.55 | -62.25 | 50 | 64, 729 | 51, 132 | 115, 861 | 32.1 | 0.06 | 0.262 | 4.074 | 0.194 | 0.758 | 0.997 | 3.702 | 5.5 | 3.07E+05 |
| St3_FL_S | 48.55 | -62.25 | 6 | 29, 398 | 63, 200 | 92, 598 | 29.3 | 19.02 | 0.71 | < 0.25 | < 0.05 | < 0.1 | < 0.5 | 0.807 | 6.3 | 2.97E+06 |
| St3_PA_CIL | 48.55 | -62.25 | 50 | 25, 280 | 23, 728 | 49, 008 | 32.1 | 0.06 | 0.262 | 4.074 | 0.194 | 0.758 | 0.997 | 3.702 | 5.5 | 3.07E+05 |
| St3_PA_S | 48.55 | -62.25 | 6 | 22, 958 | 18, 905 | 41, 863 | 29.3 | 19.02 | 0.71 | < 0.25 | < 0.05 | < 0.1 | < 0.5 | 0.807 | 6.3 | 2.97E+06 |
| St4_FL_CIL | 49.29 | -63.99 | 50 | 40, 015 | 15, 081 | 55, 096 | 32 | 0.45 | 0.181 | 3.602 | 0.212 | 0.715 | 1.656 | 2.857 | 7.4 | 4.37E+05 |
| St4_FL_S | 49.29 | -63.99 | 3 | 56, 308 | 77, 259 | 133, 567 | 28.7 | 13.9 | 1.175 | < 0.25 | < 0.05 | 0.194 | < 0.5 | 3.584 | 10 | 2.07E+06 |
| St4_PA_CIL | 49.29 | -63.99 | 50 | 40, 027 | 20, 049 | 60, 076 | 32 | 0.45 | 0.181 | 3.602 | 0.212 | 0.715 | 1.656 | 2.857 | 7.4 | 4.37E+05 |
| St4_PA_S | 49.29 | -63.99 | 3 | 17, 980 | 16, 009 | 33, 989 | 28.7 | 13.9 | 1.175 | < 0.25 | < 0.05 | 0.194 | < 0.5 | 3.584 | 10 | 2.07E+06 |
| St5_FL_CIL | 49.50 | -66.00 | 50 | 123, 718 | 53, 733 | 177, 451 | 32.2 | 0.16 | 0.064 | 6.231 | 0.252 | 0.900 | 2.214 | 6.163 | 9.7 | 2.48E+05 |
| St5_FL_S | 49.50 | -66.00 | 3 | 86, 091 | 48, 629 | 134, 720 | 29.9 | 13.6 | 0.492 | 0.273 | < 0.05 | 0.169 | < 0.5 | 1.457 | 5.1 | 1.50E+06 |
| St5_PA_CIL | 49.50 | -66.00 | 50 | 38, 300 | 15, 989 | 54, 289 | 32.2 | 0.16 | 0.064 | 6.231 | 0.252 | 0.900 | 2.214 | 6.163 | 9.7 | 2.48E+05 |
| St5_PA_S | 49.50 | -66.00 | 3 | 2, 953 | 10, 386 | 13, 339 | 29.9 | 13.6 | 0.492 | 0.273 | < 0.05 | 0.169 | < 0.5 | 1.457 | 5.1 | 1.50E+06 |
| St6_FL_CIL | 49.12 | -67.28 | 55 | 94, 924 | 44, 730 | 139, 654 | 32.4 | 0.98 | 0.075 | 12.123 | 0.113 | 1.197 | < 0.5 | 14.926 | 7.7 | 3.82E+05 |
| St6_FL_S | 49.12 | -67.28 | 3 | 30, 166 | 44, 612 | 74, 778 | 27.1 | 13.55 | 2.4314 | 1.009 | 0.058 | 0.206 | < 0.5 | 6.858 | 19.6 | 2.93E+06 |
| St6_PA_CIL | 49.12 | -67.28 | 55 | 28, 440 | 20, 979 | 49, 419 | 32.4 | 0.98 | 0.075 | 12.123 | 0.113 | 1.197 | < 0.5 | 14.926 | 7.7 | 3.82E+05 |
| St6_PA_S | 49.12 | -67.28 | 3 | 16, 001 | 8, 577 | 24, 578 | 27.1 | 13.55 | 2.4314 | 1.009 | 0.058 | 0.206 | < 0.5 | 6.858 | 19.6 | 2.93E+06 |
| St7_FL_CIL | 48.64 | -68.63 | 50 | 58, 214 | 53, 552 | 111, 766 | 32.1 | 0.74 | 0.114 | 5.385 | 0.298 | 0.853 | 2.604 | 6.414 | 16.2 | 4.38E+05 |
| St7_FL_S | 48.64 | -68.63 | 4 | 43, 382 | 48, 980 | 92, 362 | 26.9 | 8.46 | 0.481 | 7.082 | 0.140 | 0.673 | < 0.5 | 12.357 | 12.2 | 1.81E+06 |
| St7_PA_CIL | 48.64 | -68.63 | 50 | 30, 962 | 20, 055 | 51, 017 | 32.1 | 0.74 | 0.114 | 5.385 | 0.298 | 0.853 | 2.604 | 6.414 | 16.2 | 4.38E+05 |
| St7_PA_S | 48.64 | -68.63 | 4 | 19, 788 | 12, 779 | 32, 567 | 26.9 | 8.46 | 0.481 | 7.082 | 0.140 | 0.673 | < 0.5 | 12.357 | 12.2 | 1.81E+06 |

| Sample_ID | Latitude DD | Longitude DD | Depth m | Tech_rep 1 No. seqs | Tech_rep 2 No. seqs | Aggregated No. Seqs | Salinity | Temp °C | Chl A µM / ml | $NO_3^-$ µM / ml | $NO_2^-$ µM / ml | $PO_4^{3-}$ µM / ml | $NH_4^+$ µM / ml | $SiO_2^-$ µM / ml | DOC µM / ml | bacteria cells / ml |
|---|---|---|---|---|---|---|---|---|---|---|---|---|---|---|---|---|
| St8_FL_CIL | 49.98 | -66.23 | 55 | 41, 299 | 12, 562 | 53, 861 | 32.4 | 0.51 | 0.131 | 6.521 | 0.252 | 0.925 | 1.518 | 6.916 | 4.7 | 3.14E+05 |
| St8_FL_S | 49.98 | -66.23 | 5 | 71, 411 | 56, 127 | 127, 538 | 29.5 | 15.04 | 0.7 | < 0.25 | < 0.05 | 0.188 | < 0.5 | 2.032 | 13.3 | 1.52E+06 |
| St8_PA_CIL | 49.98 | -66.23 | 55 | 32, 121 | 26, 153 | 58, 274 | 32.4 | 0.51 | 0.131 | 6.521 | 0.252 | 0.925 | 1.518 | 6.916 | 4.7 | 3.14E+05 |
| St8_PA_S | 49.98 | -66.23 | 5 | 24, 016 | 31, 562 | 55, 578 | 29.5 | 15.04 | 0.7 | < 0.25 | < 0.05 | 0.188 | < 0.5 | 2.032 | 13.3 | 1.52E+06 |
| St10_FL_CIL | 50.22 | -58.47 | 65 | 47, 535 | 44, 129 | 91, 664 | 32.3 | -0.1 | 0.065 | 8.693 | 0.183 | 1.065 | 0.813 | 10.498 | 5.9 | 2.86E+05 |
| St10_FL_S | 50.22 | -58.47 | 3 | 51, 873 | 25, 686 | 77, 559 | 30.4 | 16.66 | 0.625 | < 0.25 | < 0.05 | 0.199 | < 0.5 | 3.633 | 15.5 | 1.30E+06 |
| St10_PA_CIL | 50.22 | -58.47 | 65 | 40, 809 | 37, 384 | 78, 193 | 32.3 | -0.1 | 0.065 | 8.693 | 0.183 | 1.065 | 0.813 | 10.498 | 5.9 | 2.86E+05 |
| St10_PA_S | 50.22 | -58.47 | 3 | 9, 844 | 7, 423 | 17, 267 | 30.4 | 16.66 | 0.625 | < 0.25 | < 0.05 | 0.199 | < 0.5 | 3.633 | 15.5 | 1.30E+06 |

**Supplementary Table S2** Wilcoxon test comparing the mean relative abundances of phyla between free-living (FL) and particle-attached (PA) lifestyles at surface and cold-intermediate layer (CIL). 'Unidentified_Bacteria' are a combination of OTUs unable to be assigned taxonomy at the phylum level. 'Others' are all OTUs assigned phyla that collectively (across samples) comprise less than 0.01 relative abundance of the total reads. This value is based on the pairwise difference between the individuals in your two groups. "V" value is based on pairwise difference between the individuals in the two groups, following a certain probability distribution. Bold values are significant at $p < 0.05$.

| Free-living vs. Particle-associated lifestyles | | | | |
|---|---|---|---|---|
| Phylum | Surface | | CIL | |
| | V | p-value | V | p-value |
| Acidobacteria | 36 | 0.293 | 34.5 | 0.282 |
| Actinobacteria | 75 | **0.001** | 51 | **0.193** |
| BD1-5 | 39.5 | 0.48 | 40 | 0.5 |
| Unclassified | 13 | **0.007** | 14 | **0.009** |
| Bacteroidetes | 20 | **0.039** | 54 | 0.889 |
| Candidate_BRC1 | 40 | 0.5 | 26 | 0.107 |
| Candidate_division_OD1 | 41.5 | 0.56 | 31 | 0.213 |
| Candidate_TM7 | 28.5 | 0.15 | 23 | 0.065 |
| Chlamydiae | 26 | 0.107 | 11 | **0.004** |
| Chloroflexi | 31.5 | 0.186 | 61 | **0.039** |
| Cyanobacteria | 17 | **0.02** | 19 | **0.031** |
| Deferribacteres | 62 | **0.032** | 69 | **0.007** |
| Firmicutes | 7 | **0.002** | 5.5 | **0.001** |
| Fusobacteria | 34 | 0.292 | 35 | 0.322 |
| Gemmatimonadetes | 26 | 0.059 | 22.5 | 0.061 |
| Lentisphaerae | 25 | **0.09** | 10 | 0.003 |
| Planctomycetes | 4 | **0.014** | 4 | 0.014 |
| Alphaproteobacteria | 67 | **0.009** | 74 | **0.001** |
| Betaproteobacteria | 66 | **0.012** | 69 | **0.005** |
| Deltaproteobacteria | 11 | **0.004** | 16 | **0.016** |
| Epsilonproteobacteria | 38.5 | 0.447 | 21 | **0.047** |
| Gammaproteobacteria | 52 | 0.17 | 28 | 0.149 |
| TM6 | 34.5 | 0.297 | 26 | **0.08** |
| Tenericutes | 25.5 | 0.523 | 32 | 0.224 |
| Verrucomicrobia | 12 | **0.005** | 29 | 0.17 |
| Others | 38.5 | 0.428 | 22.5 | 0.058 |

**Supplementary Table S3** The proportion of specialists and generalists in terms of the number of taxa at different taxonomical levels for the individual sub-communities defined by depth (surface or cold intermediate layer: CIL) and size fractionation (free-living: FL or particle-attached: PA). 'Prop. Spec %' and 'Prop. Gener %' refer to the proportion of specialists and generalists in the respective community, respectively.

| Taxonomic rank | Surface | | | |
|---|---|---|---|---|
| | FL | | PA | |
| | Prop. Spec % | Prop. Gener % | Prop. Spec % | Prop. Gener % |
| Species (OTU 99%) | 83.96 | 3.06 | 87.74 | 3.63 |
| Genus | 21.28 | 18.96 | 37.11 | 20.87 |
| Family | 20.06 | 27.12 | 30.79 | 28.81 |
| Order | 18.08 | 30.51 | 29.38 | 35.03 |
| Class | 21.28 | 24.47 | 35.11 | 29.79 |
| Phylum | 17.24 | 31.03 | 20.69 | 44.83 |
| | CIL | | | |
| | FL | | PA | |
| | Prop. Spec % | Prop. Gener % | Prop. Spec % | Prop.gener % |
| Species (OTU 99%) | 75.04 | 4.45 | 79.89 | 7.01 |
| Genus | 27.01 | 27.01 | 22.51 | 37.11 |
| Family | 21.47 | 37.85 | 18.36 | 47.18 |
| Order | 19.21 | 44.07 | 16.38 | 51.98 |
| Class | 18.09 | 41.49 | 18.09 | 51.06 |
| Phylum | 3.45 | 55.17 | 6.9 | 72.41 |

**Supplementary Table S4** The relative abundance of specialists **A)** and generalists **B)** at the class-level for the individual sub-communities defined by depth (surface or cold intermediate layer: CIL) and size-fractionation (free-living: FL or particle-attached: PA). 'SUM' in the last row of the tables represent the sum of relative abundances of all taxa (in a column).

**A)** Specialist

| Specialist_class | Surface_FL | Surface_PA | CIL_FL | CIL_PA |
|---|---|---|---|---|
| 028H05-P-BN-P5(100) | 0 | 0.000333192 | 0.00041649 | 0 |
| Acidobacteria_unclassified(100) | 0 | 0 | 0 | 0.000333192 |
| Acidobacteria(100) | 0 | 0.007413516 | 0.000749681 | 0 |
| Actinobacteria_unclassified(100) | 8.33E-05 | 0 | 0 | 0 |
| AEGEAN-245(100) | 0.00041649 | 0.000333192 | 0 | 0 |
| A0erolineae(100) | 0 | 0.003331917 | 0.00041649 | 0 |
| Arctic97B-4_marine_group(100) | 0.002249044 | 0.000749681 | 0 | 0 |
| Ardenticatenia(100) | 0 | 0.000333192 | 0 | 0.001166171 |
| Armatimo0detes_unclassified(100) | 0.000166596 | 0.006580537 | 0 | 0 |
| Bacteroidetes_Incertae_Sedis(100) | 0.000166596 | 0.001082873 | 0 | 0 |
| Bacteroidia(100) | 8.33E-05 | 0.002748832 | 0.00041649 | 0 |
| BD2-2(100) | 0 | 0.001332767 | 0 | 0 |
| BD7-11(100) | 0 | 0 | 8.33E-05 | 0.001665959 |
| Caldilineae(100) | 0 | 0.002082448 | 0 | 0 |
| Candidate_division_BRC1_unclassified(100) | 0.000499788 | 0.005331068 | 0 | 0 |
| Candidate_division_SR1_unclassified(100) | 0 | 0.000916277 | 0 | 0 |
| Candidate_division_WS3_unclassified(100) | 0 | 0 | 0 | 0.000499788 |
| Chlorobia(100) | 0 | 0.000916277 | 0 | 0.000249894 |
| Chloroflexi_unclassified(100) | 0 | 0.000333192 | 0 | 0.000749681 |
| Clostridia(100) | 0.000166596 | 0 | 0 | 0 |
| Dehalococcoidia(100) | 0 | 0 | 0 | 0.000583086 |
| DEV055(100) | 0 | 0.001665959 | 0 | 0.000749681 |
| Erysipelotrichia(100) | 0 | 0.000333192 | 0.000166596 | 0.000499788 |
| Fibrobacteria(100) | 0 | 0.001249469 | 0 | 0 |
| Gemmatimo0detes(100) | 0.00041649 | 0 | 0 | 0 |
| Holophagae(100) | 0 | 0.000583086 | 0 | 0 |
| Ig0vibacteria(100) | 0 | 0.000832979 | 0 | 0 |

| Specialist_class | Surface_FL | Surface_PA | CIL_FL | CIL_PA |
|---|---|---|---|---|
| JG30-KF-CM66(100) | 8.33E-05 | 0 | 0.001416065 | 0 |
| JTB23(100) | 0.000499788 | 0 | 0 | 0 |
| Ktedonobacteria(100) | 0 | 0.000916277 | 0 | 0 |
| Lentisphaeria(100) | 0.00241564 | 0 | 0 | 0 |
| MB-A2-108(100) | 0 | 0.001249469 | 0 | 0.000249894 |
| Melai0bacteria(100) | 0 | 0.001416065 | 0 | 0 |
| ML635J-21(100) | 0.001166171 | 0 | 0 | 0 |
| Mollicutes(100) | 8.33E-05 | 0 | 0 | 0 |
| Negativicutes(100) | 0 | 0.000333192 | 0 | 0.000666383 |
| Oligosphaeria(100) | 0 | 0.004997876 | 0 | 0 |
| OPB35_soil_group(100) | 0.000832979 | 0 | 0 | 0 |
| OPB54(100) | 0 | 0 | 0 | 0.000583086 |
| PBS-III-20(100) | 0 | 0.00041649 | 0 | 0 |
| Pla3_lineage(100) | 0.001416065 | 0 | 0 | 0 |
| Planctomycetes_unclassified(100) | 0.000916277 | 0 | 0.002249044 | 0 |
| Proteobacteria_Incertae_Sedis(100) | 0 | 0.000916277 | 0 | 0.00041649 |
| R76-B128(100) | 0.000166596 | 0 | 8.33E-05 | 0 |
| SAR202_clade(100) | 0.006330643 | 0.00241564 | 0 | 0 |
| SGST604(100) | 0 | 0 | 0.001166171 | 0 |
| SL56_marine_group(100) | 0 | 0.00199915 | 0.001332767 | 0.000832979 |
| Spartobacteria(100) | 0 | 0 | 0.003082023 | 0 |
| SPG12-401-411-B72(100) | 0 | 0 | 0.000249894 | 0.004248194 |
| SS1-B-03-39(100) | 0 | 0.005497664 | 0.001082873 | 0 |
| Subgroup_22(100) | 0 | 0 | 0 | 0.000583085 |
| TM6_unclassified(100) | 0 | 0 | 0.000249894 | 0 |
| VC2.1_Bac22(100) | 0 | 0.000583086 | 0 | 0 |
| Verrucomicrobia_Incertae_Sedis(100) | 0 | 0 | 0.001915852 | 0.000166596 |
| Verrucomicrobia_unclassified(100) | 0.001582661 | 0 | 0 | 0 |
| WCHB1-41(100) | 0 | 0.000916277 | 0.000249894 | 0 |
| WCHB1-60_unclassified(100) | 0 | 0.001499363 | 0 | 0 |
| SUM | 0.01974161 | 0.06164047 | 0.015326819 | 0.014243946 |

**B)** Generalists

| Generalist_class | Surface_FL | Surface_PA | CIL_FL | CIL_PA |
|---|---|---|---|---|
| Acidimicrobiia(100) | 5.284170894 | 2.084364145 | 2.030387085 | 1.511107779 |
| Actinobacteria(100) | 0.825898993 | 1.187162123 | 0.482961408 | 0.728273817 |
| AEGEAN-245(100) | 0 | 0 | 0.017825757 | 0 |
| Alphaproteobacteria(100) | 44.99979176 | 14.56964124 | 36.0935769 | 7.592939668 |
| Arctic97B-4_marine_group(100) | 0 | 0 | 0.128945198 | 0 |
| Bacilli(100) | 0 | 0.054310251 | 0.014327244 | 0.118699553 |
| Bacteria_unclassified(100) | 0.054976635 | 2.063456364 | 0.234733572 | 1.090120032 |
| Bacteroidetes_unclassified(100) | 0.439229994 | 3.889930113 | 0.368843242 | 2.099357773 |
| Bacteroidia(100) | 0 | 0 | 0 | 0.007746708 |
| BD1-5_unclassified(100) | 0 | 0 | 0.020407993 | 0.029654064 |
| BD2-2(100) | 0 | 0 | 0 | 0.00924607 |
| Betaproteobacteria(100) | 1.400071636 | 0.903699261 | 1.363920334 | 0.840809323 |
| Candidate_division_BRC1_unclassified(100) | 0 | 0 | 0.004914578 | 0.079882716 |
| Candidate_division_OD1_unclassified(100) | 0 | 0 | 0.023073527 | 0.03181981 |
| Candidate_division_SR1_unclassified(100) | 0 | 0 | 0 | 0.008912879 |
| Candidate_division_TM7_unclassified(100) | 0 | 0.10953678 | 0 | 0.224987713 |
| Chlamydiae(100) | 0 | 0.454723409 | 0.045314075 | 0.382087613 |
| Clostridia(100) | 0 | 0.034901833 | 0 | 0.028071403 |
| Cyanobacteria_unclassified(100) | 0.020074802 | 0.445810531 | 0.003415215 | 0.071636221 |
| Cyanobacteria(100) | 7.18727874 | 11.91443636 | 0.307369368 | 2.794895503 |
| Cytophagia(100) | 0.505118658 | 0.258806674 | 0.551598904 | 0.568924874 |
| Deferribacteres(100) | 0.126779452 | 0.022990229 | 0.74226787 | 0.030903533 |
| Deltaproteobacteria(100) | 0.607908306 | 2.381321272 | 2.617470908 | 5.939725616 |
| Epsilonproteobacteria(100) | 0.026322146 | 0.075384628 | 0.106288161 | 0.266969871 |
| Flavobacteriia(100) | 18.5661094 | 25.36005531 | 17.40401996 | 14.49425661 |
| Fusobacteriia(100) | 0 | 0.033569066 | 0 | 0.019991504 |
| Gammaproteobacteria(100) | 14.6678495 | 10.1932512 | 25.97462745 | 27.56228603 |

| Generalist_ class | Surface_FL | Surface_PA | CIL_FL | CIL_PA |
|---|---|---|---|---|
| Gemmatimo0detes(100) | 0 | 0 | 0.019075226 | 0.068804092 |
| Lentisphaerae_unclassified(100) | 0 | 0.124613706 | 0.005414365 | 0.384753147 |
| Lentisphaeria(100) | 0 | 0 | 0.064222705 | 0.603576813 |
| ML635J-21(100) | 0 | 0 | 0.020241397 | 0.018242247 |
| Mollicutes(100) | 0 | 0 | 0 | 0.004248195 |
| Oligosphaeria(100) | 0 | 0 | 0.01690948 | 0.21265962 |
| OM190(100) | 0.02457289 | 3.206720478 | 0.195583544 | 1.688449076 |
| OPB35_soil_group(100) | 0 | 0 | 0.021157675 | 0.012911179 |
| Opitutae(100) | 0.696787199 | 1.937509892 | 0.344187054 | 0.390417406 |
| Phycisphaerae(100) | 0.059974511 | 0.891787657 | 0.686291659 | 4.369976094 |
| Pla3_lineage(100) | 0 | 0 | 0.02257374 | 0.135442437 |
| Planctomycetacia(100) | 0.394082515 | 6.446593531 | 1.620644559 | 14.32249627 |
| Planctomycetes_unclassified(100) | 0 | 0 | 0 | 0.0443145 |
| Proteobacteria_unclassified(100) | 0.054560145 | 0.098957943 | 0.138607758 | 0.545934645 |
| SAR202_clade(100) | 0 | 0 | 0.300205746 | 0.010412241 |
| SC3-20(100) | 0.042981733 | 0 | 0.02940417 | 0 |
| Spartobacteria(100) | 0 | 0 | 0 | 0.011911604 |
| Sphingobacteriia(100) | 0.456472666 | 1.358755862 | 0.536521978 | 0.812071536 |
| SPOTSOCT00m83(100) | 0.095292834 | 0.023906506 | 0.201997485 | 0.014243946 |
| SS1-B-03-39(100) | 0 | 0 | 0 | 0.006913728 |
| Thermoleophilia(100) | 0 | 0 | 0 | 0.009079475 |
| TM6_unclassified(100) | 0 | 0 | 0 | 0.008829581 |
| Verrucomicrobia_unclassified(100) | 0 | 0 | 0.07854995 | 0.010328943 |
| Verrucomicrobiae(100) | 3.41479871 | 9.605417698 | 7.122972736 | 9.66289327 |
| SUM | 99.95110412 | 99.73161406 | 99.96084997 | 99.88221672 |

**Supplementary Table S5** The slope of the abundance-occupancy relationships along the taxonomic ranks **A)** and fold change of niche index between consecutive taxonomic ranks **B)** for the individual sub-communities defined by depth (surface or cold intermediate layer: CIL) and size fractionation (free-living: FL or particle-attached: PA). For each rank pair in **B)**, the value obtained from the low taxonomic rank was used as a reference value to that from the high rank.

**A)**

| | *Slope of AORs* | | | |
|---|---|---|---|---|
| Ranks | Surface_FL | Surface_PA | CIL_FL | CIL_PA |
| Species | 0.69 | 0.81 | 0.69 | 0.65 |
| Genus | 0.84 | 0.72 | 0.79 | 0.70 |
| Family | 0.81 | 0.73 | 0.80 | 0.72 |
| Order | 0.83 | 0.78 | 0.85 | 0.79 |
| Class | 0.97 | 0.90 | 0.89 | 0.91 |
| Phylum | 1.15 | 0.95 | 1.05 | 1.06 |

**B)**

| | *Fold change* | | | |
|---|---|---|---|---|
| Rank pairs | Surface_FL | Surface_PA | CIL_FL | CIL_PA |
| Species -> Genus | 0.217 | -0.111 | 1.145 | 0.077 |
| Genus -> Family | -0.036 | 0.014 | 0.013 | 0.029 |
| Family -> Order | 0.025 | 0.068 | 0.063 | 0.097 |
| Order -> Class | 0.169 | 0.154 | 0.047 | 0.152 |
| Class -> Phylum | 0.186 | 0.056 | 0.180 | 0.165 |

### Supplementary Figures

**Supplementary Figure S1** Non-metric multidimensional scaling (NMDS) based on Morista-Horn and generalized UniFrac matrices illustrating the beta-diversity of the particle-associated (triangles) and free-living bacteria (circles) in the surface water (orange) and cold intermediate layer (dark olive green). Filled and unfilled symbols refer to technical sequencing replicates. The Horn-Morisita distance metric (Horn, 1996) was chosen to calculate beta diversity because it is more robust to uneven samples (i.e., no normalization of the dataset) than the commonly used Bray-Curtis distance. Generalized UniFrac distance was used to compromise the biases from the unweighted and weighted UniFrac matrices, so that the distance was not dominated by highly abundant lineages (R package 'GUniFrac'; Chen et al., 2012). Closely clustered symbols indicate bacterial communities that were more similar than those further apart from each other in the multidimensional ordination.

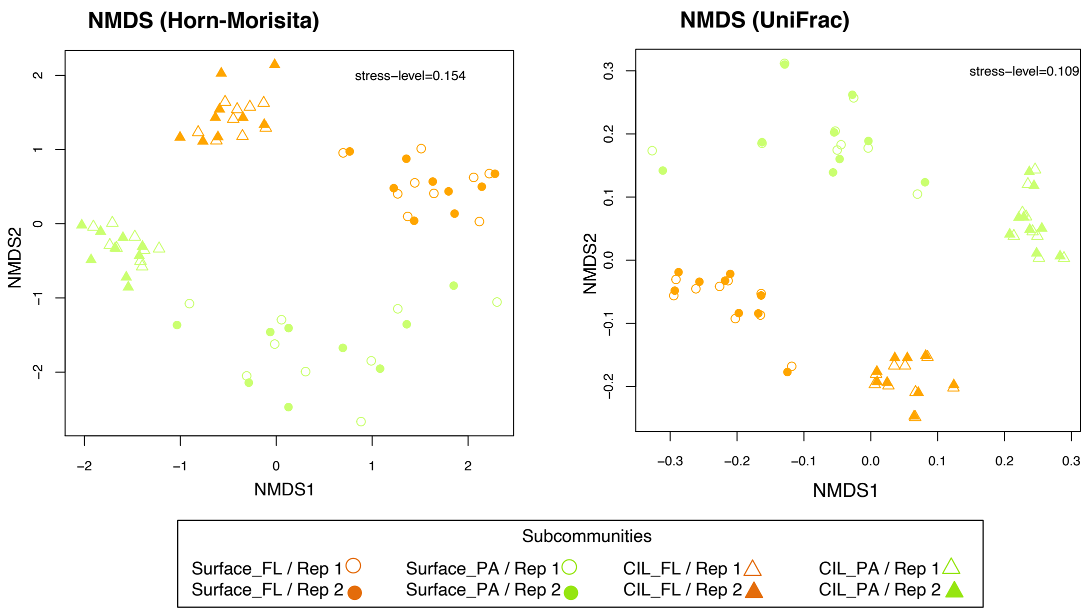

**Supplementary Figure S2** Non-metric multidimensional scaling (NMDS) based on Morista-Horn illustrating the beta-diversity of the particle-associated (triangles) and free-living bacteria (circles) in the surface water (orange) and cold intermediate layer (dark olive green). Filled and unfilled symbols refer to technical sequencing replicates. The analysis was done using the exact sequence variants generated from DADA2 pipeline (Callahan et al., 2017). Raw sequences were processed using the DADA2 pipeline (Callahan et al., 2017) according to the DADA2 tutorial (v1.12) in R. The sequences were quality filtered with customized modifications as follows: truncLen=c(285,205), maxEE=4, truncQ=2, maxN=0, rm.phix=TRUE, trimLeft=c(17,22). Subsequently, denoising, merging and chimera removal were completed according to the DADA2 pipeline tutorial. All sequences were aligned and assigned taxonomically using the SILVA v.123 reference database. All archaea, eukaryote, mitochondria, and chloroplast sequences were removed. Singletons (ASVs with only one sequence read across all samples) were also discarded.

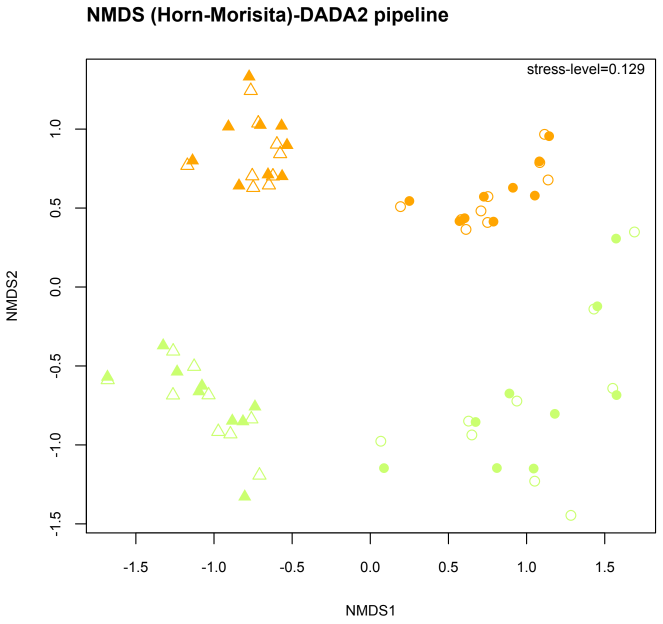

**Supplementary Figure S3** Occupancy (number of sites) plotted against the mean relative abundance across taxonomic ranks from the species to the phylum level (**A–E,** except for the class level which is presented in Figure 4). The number of the occupied sites was used to define the niche breadth of bacteria. Taxa that occupied fewer than two stations were defined as habitat specialists (red), and those occupying more than five stations as habitat generalists. The proportion of the two ecological groups is shown on the figure. Correlation coefficient and *P* value obtained from Spearman colerration analysis are presented on the lower right corner of each panel.

**A. Species level (OTU-level at 99% sequence similarity)**

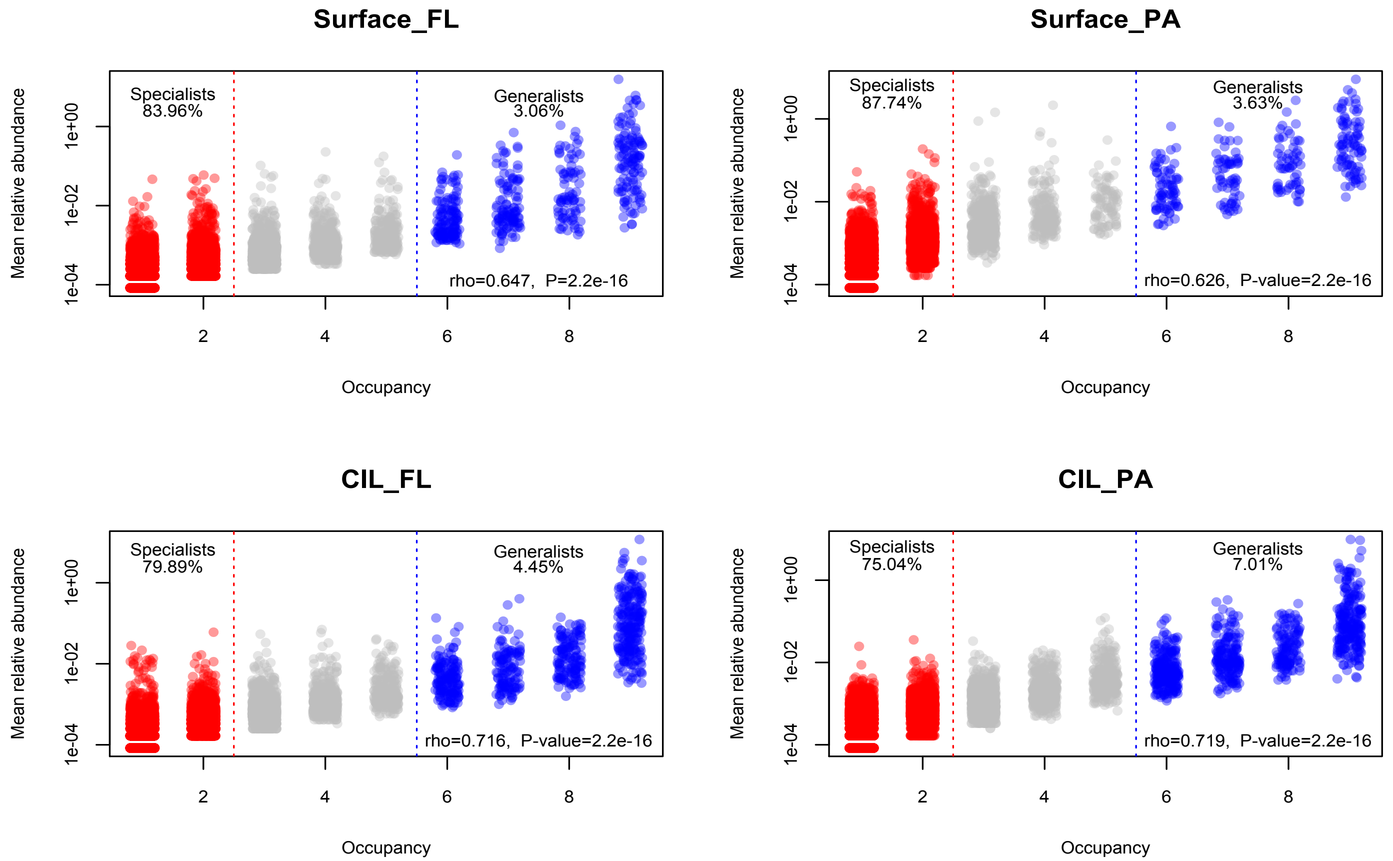

**B. Genus level**

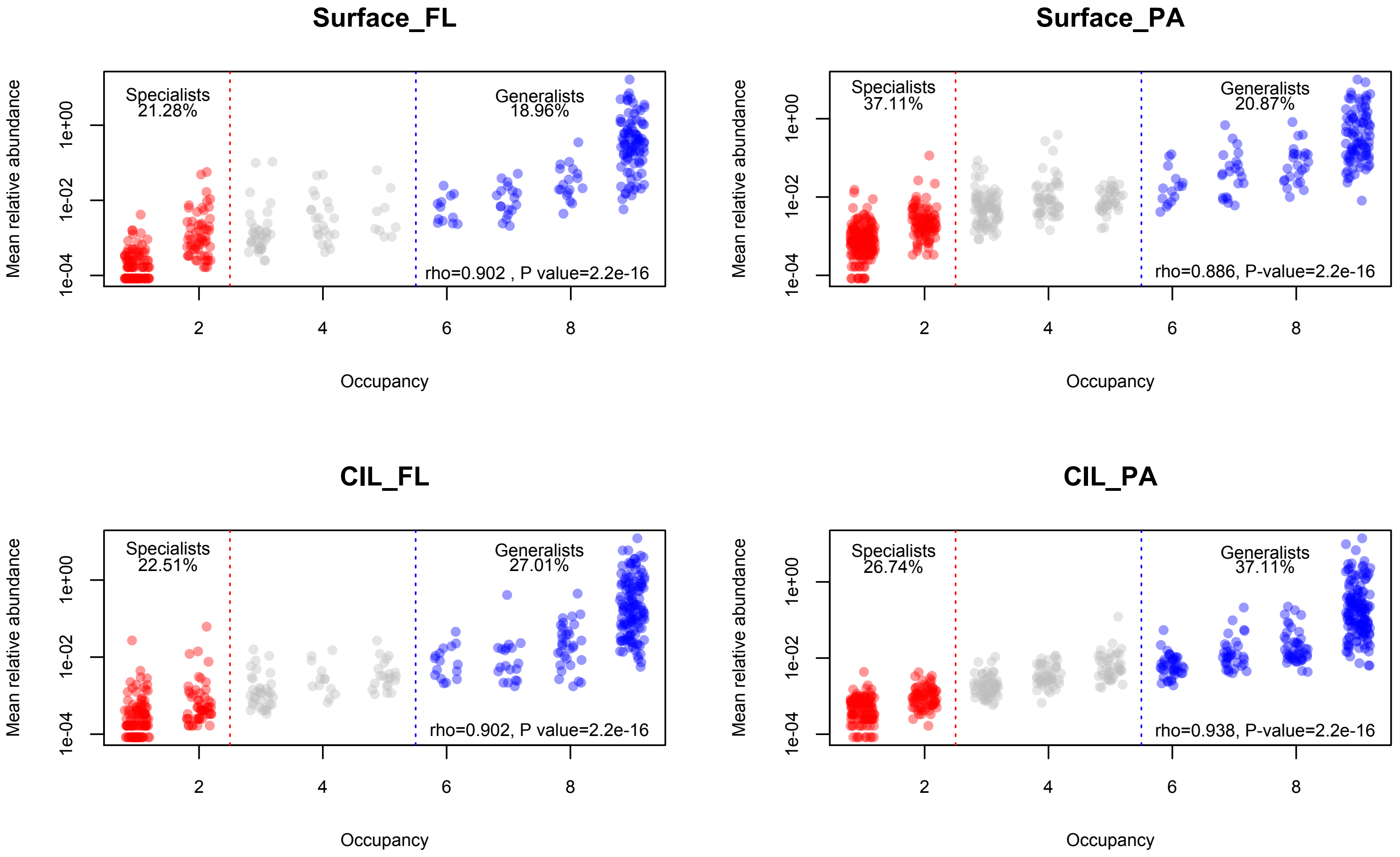

### C. Family level

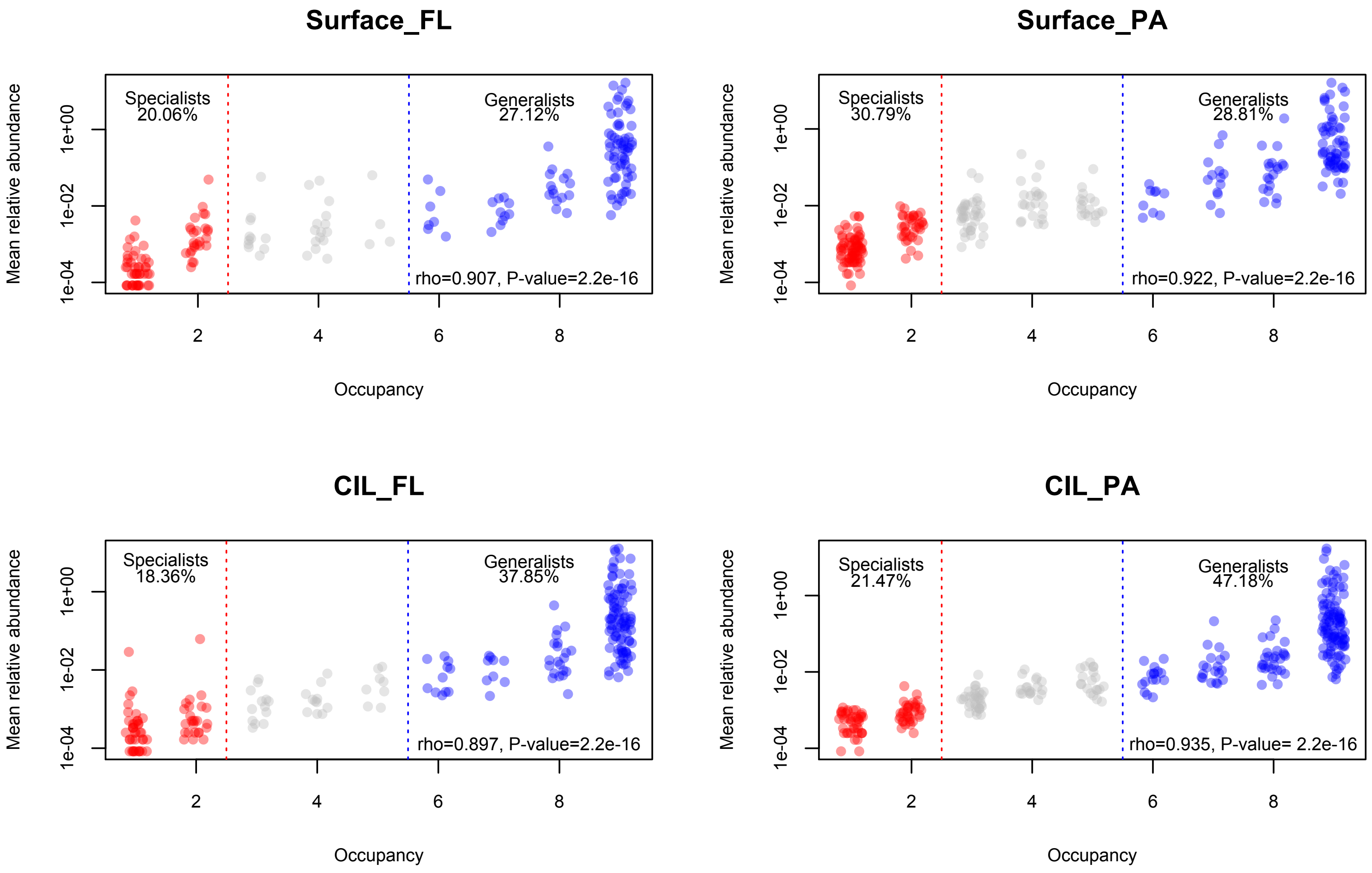

**D. Order level**

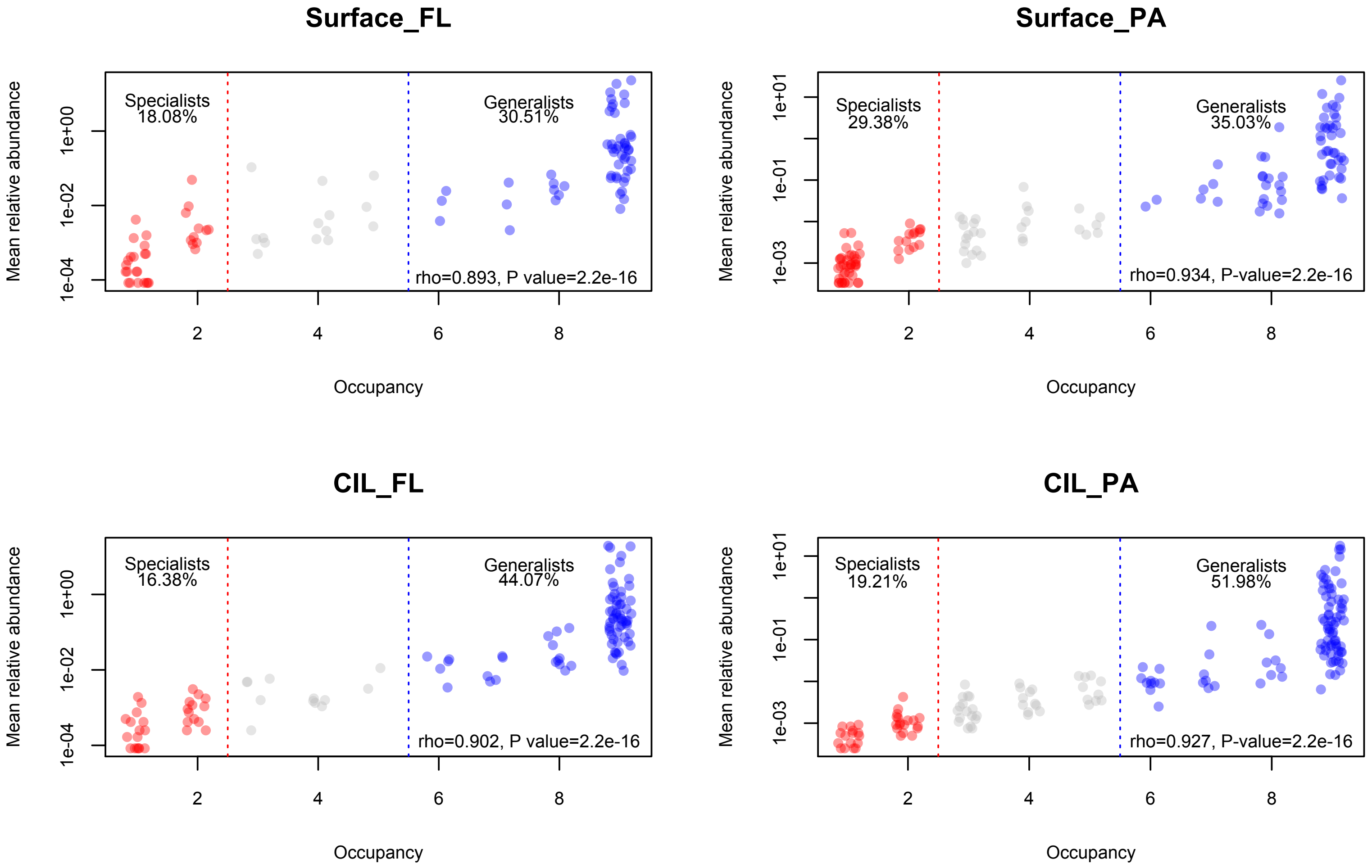

### E. Phylum level

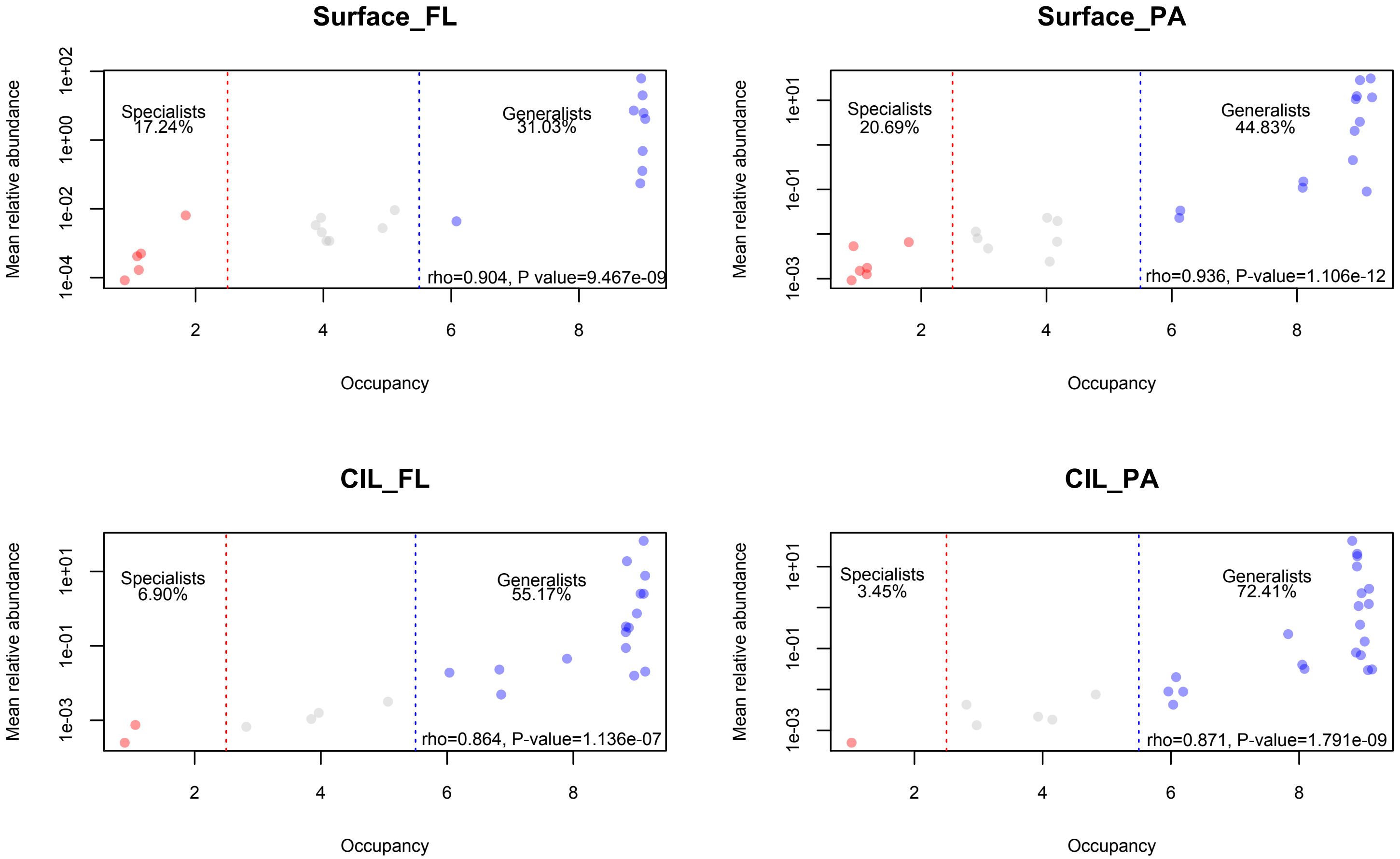

**Supplementary Figure S4** Relative proportion of the top 20 bacterial classes among habitat specialists and generalists. For either ecological group, the proportion within each group was calculated from the percentage of the relative abundance of a given class / the aggregated relative abundance of the respective sample. The relative abundance of habitat specialists and generalists at the class level are given in Supplementary Table S4.

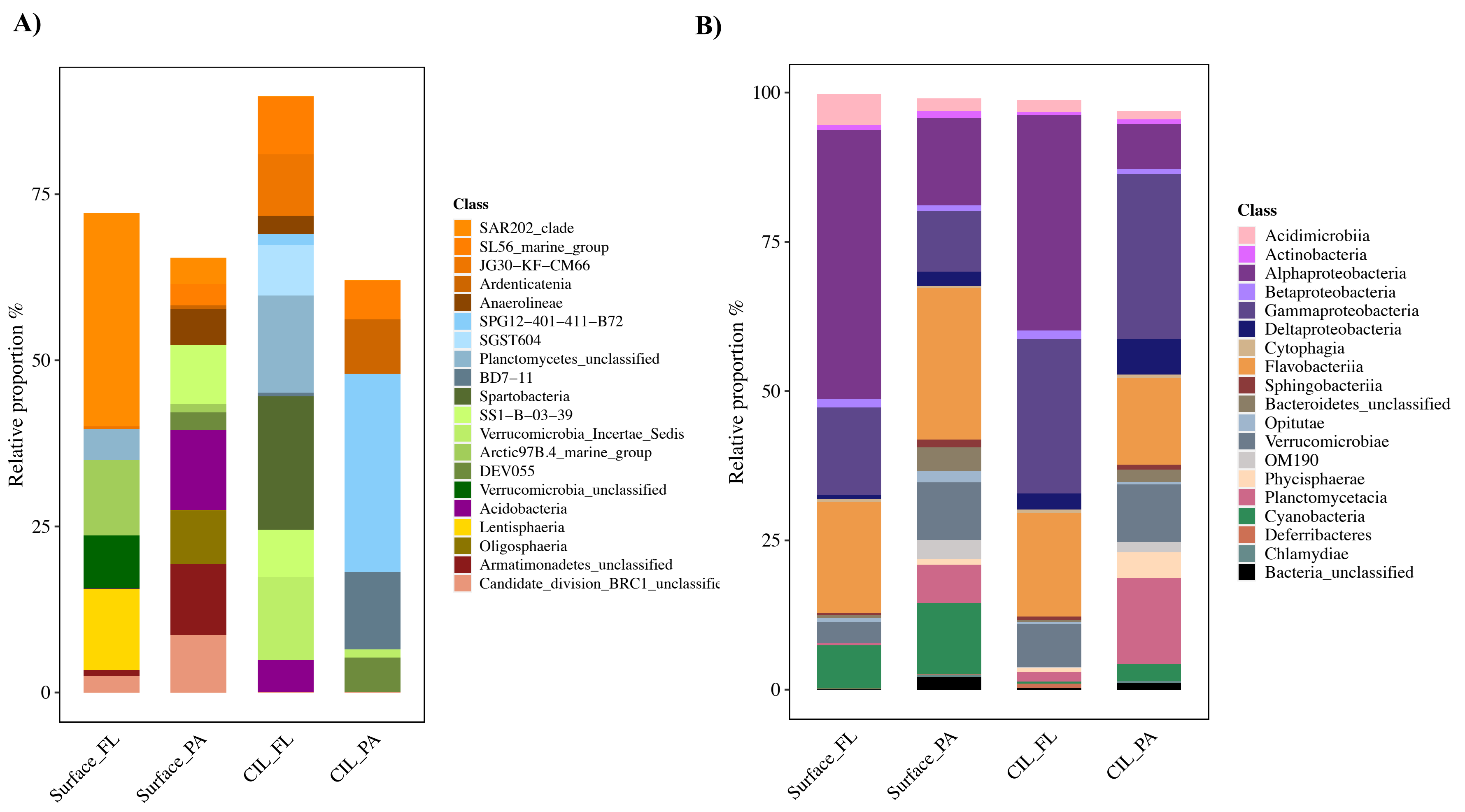
